# Signatures of replication timing, recombination and sex in the spectrum of rare variants on the human X chromosome and autosomes

**DOI:** 10.1101/519421

**Authors:** Ipsita Agarwal, Molly Przeworski

## Abstract

The sources of human germline mutations are poorly understood. Part of the difficulty is that mutations occur very rarely, and so direct pedigree-based approaches remain limited in the numbers that they can examine. To address this problem, we consider the spectrum of low frequency variants in a dataset (gnomAD) of 13,860 human X chromosomes and autosomes. X-autosome differences are reflective of germline sex differences, and have been used extensively to learn about male versus female mutational processes; what is less appreciated is that they also reflect chromosome-level biochemical features that differ between the X and autosomes. We tease these components apart by comparing the mutation spectrum in multiple genomic compartments on the autosomes and between the X and autosomes. In so doing, we are able to ascribe specific mutation patterns to replication timing and recombination, and to identify differences in the types of mutations that accrue in males and females. In particular, we identify C>G as a mutagenic signature of male meiotic double strand breaks on the X, which may result from late repair. Our results show how biochemical processes of damage and repair in the germline interact with sex-specific life history traits to shape mutation patterns on both the X chromosome and autosomes.

## Introduction

Germline mutations, the source of all heritable variation, accrue each generation from accidental changes to the genome during the development of gametes. These mutations reflect a balance of exogenous damage or endogenous processes that alter DNA in the germline, and processes that correctly repair DNA lesions before the next replication (1). The biochemical machinery that underlies germline mutagenesis can be conceptualized as a set of genetic loci that modulate the net mutational input in each generation, and variants in these loci as “modifiers of mutation” (2, 3). Since the activity of distinct biochemical pathways often leaves different signatures in DNA (4–10), these modifiers influence the distribution of mutation types (the “mutation spectrum”), as well as the total number of mutations inherited by offspring.

The mutational landscape in the germline is also modified by the sex of the parent: in humans, notably, it has long been known that males contribute three times as many mutations on average as females per generation (11, 12). As in other mammals, gametogenesis differs drastically by sex: female germ cells are arrested in meiosis for much of their development whereas male germ cells enter meiosis late in their development (13–15). Male germ cells undergo many more cell divisions than female germ cells; they are also methylated earlier and have higher methylation levels on average throughout ontogenesis (16). Due to differences in their cellular biochemistry at different developmental stages, male and female gametes may be subject to different kinds of endogenous and environmental insults, or repair different types of DNA lesions with varying degrees of efficacy. For example, male gametes may accrue oxidative damage due to lack of base excision repair in late spermatogenesis (17). Males and females also differ in life history traits such as the timing of puberty and age of reproduction (18), which modulate the exposure of the gamete to the biochemical states associated with particular stages of development and thus alter their mutagenic impact. In that sense, the sex of the parent as well as variants in loci associated with sex-specific biochemistry and life history are also modifiers of mutation. The germline mutation spectrum in each generation is therefore a convolution of the signatures left by biochemical machinery in DNA sequence and the influence of sex on the developmental trajectories of germ cells.

In principle, it is possible to characterize mutational mechanisms by decomposing the mutation spectrum into its component signatures. Such an approach has led to a wealth of insight into the sources of somatic mutations, i.e., mutations that accumulate in somatic tissues during normal development or ageing. Distinct signatures of processes that generate or repair DNA lesions have been identified by analyzing millions of somatic mutations in their immediate sequence context, across tumor samples of diverse etiologies (4, 5, 7, 10). A complementary approach, based on changes in the mutation spectrum with regional variation in genomic features, has further illuminated the influence of local replication timing, transcription, chromatin organization, and epigenetic modifications on somatic mutagenesis (19–28).

These methods have proved difficult to apply to the germline however, because each offspring inherits only about 70 de novo mutations on average (29). Thus, the most direct approach to the study of germline mutations, the resequencing of pedigrees (12, 29–32), remains limited in its ability to identify determinants of mutation rate variation. For instance, examining 96 possible mutation types considered in a trinucleotide context in ∼100,000 de novo mutations, the biggest study to date found only three mutation types for which the proportion transmitted from mothers and fathers differed significantly (29). Additionally, the mutation patterns from the three largest de novo mutation studies combined show inconsistent patterns of correlation to genomic features, for reasons that remain unclear (33).

One way to overcome the limitation of small samples in studies of germline mutation is to use polymorphisms as a proxy for de novo mutations. In large samples, most polymorphisms are rare and recent enough for effects of direct and indirect selection and biased gene conversion to be minimal; they should therefore recapitulate the de novo mutation spectrum with reasonable fidelity (31, 34, 35). The much higher density of rare variants across the genome can then be used to more robustly investigate associations with genomic features. Using this strategy, a recent study of human autosomal data identified mutation types and contexts significantly associated with a variety of genomic features (34). While the authors suggested putative biochemical sources for three signatures in the germline based on their similarity to patterns that have been reported in tumors, it is unclear to what degree these mechanisms can be directly extrapolated to the germline (36, 37). Moreover, sex-specific effects on the mutation spectrum were not considered.

Insight into sex-specific effects can be gained by contrasting polymorphism levels on the sex chromosomes and autosomes, since autosomes reflect mutational processes in the male and female germlines equally, while the X chromosome disproportionately reflects the female germline, and the Y chromosome exclusively reflects the male germline. This approach to studying sex differences has been used extensively; notably, its application to divergence data provided the first systematic evidence for a higher contribution of males to mutation in humans and other mammals (38, 39). Yet no significant influence of sex on the mutation spectrum was inferred in a recent comparison of ∼3000 rare variants on the X and Y chromosomes (31). Despite their importance, therefore, the genesis of germline mutations remains poorly understood to date, and the role of sex-specific modifiers particularly enigmatic.

To fill this gap, we consider the spectrum of rare polymorphisms across the genome using genome-wide SNPs in the gnomAD dataset (40). We compare particular genomic “compartments”, or units of the genome with unique combinations of biochemical and sex-specific properties, on the X and autosomes; this approach enables us to tease apart biochemical and sex-specific influences on the germline mutation spectrum. With over 120 million SNPs to analyze across the genome, we can thus detect even subtle differences in mutational patterns between genomic compartments.

## Materials and Methods

We use whole genome SNP data from 15,496 individuals made available by the Genome Aggregation Database (gnomAD), which includes 9,256 Europeans and 4,368 African or African-American individuals (40). We limit our analysis to the 6,930 female individuals in the dataset to sample X-chromosomes and autosomes in equal numbers. We then compare the diversity levels of different mutation types in pairs of genomic compartments (Fig. 1a). In these data, there are ∼120 million SNPs, of which 53% of the variants are singletons (i.e., variants seen only once in the sample, with an allele count of one), and 11% are doubletons (allele count = 2) (Fig. 1b). Only about 10% of variants are at frequency 1% or greater; we retain them, given that their inclusion does not affect our qualitative results (SI Appendix, Fig. S1).

**Fig. 1.**
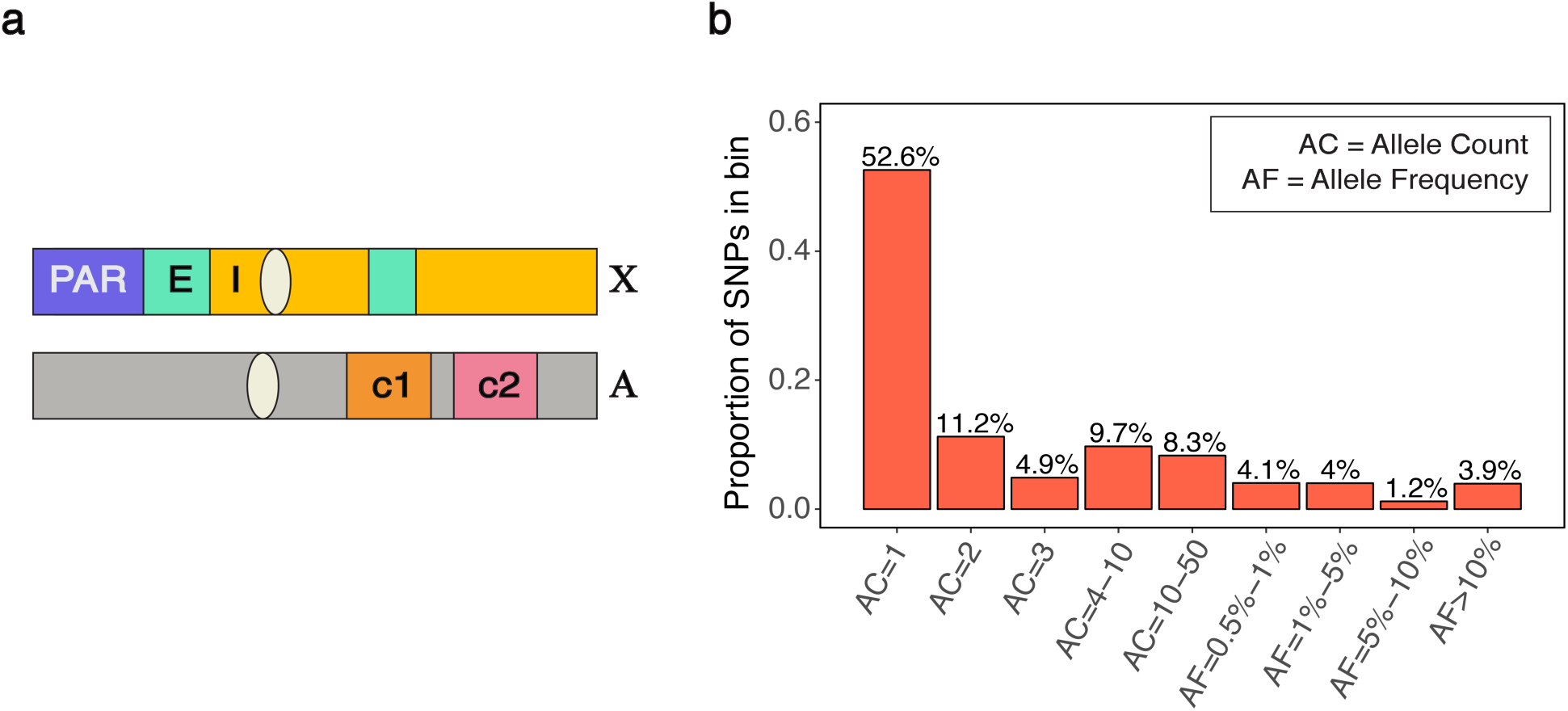
(a) A schematic of genomic compartments on the X chromosome and autosomes. Three compartments on the X chromosome are depicted: the pseudoautosomal region (PAR), regions of the X that escape inactivation (E), regions of the X that undergo inactivation (I). Also depicted are two hypothetical autosomal compartments with distinct biochemical properties (c1 and c2). Analyses include pairwise comparisons of mutational patterns between autosomal compartments and between the X chromosome and autosomes. (b) The frequency spectrum of variants (N = 120,521,915) in the 13,860 chromosomes analyzed. Over two-thirds of mutations are present at three copies or less in the sample.

As in other recent studies, we extract the single base pair flanking sequence on each side of the variant position using the hg19 reference to obtain mutations in their trinucleotide context, and combine mutations in reverse complement classes (for example, the ACG>ATG and CGT>CAT classes are collapsed into the former) to obtain 96 mutation types. Unless otherwise noted, we treat the major allele as the ancestral state at a site; however, we obtain similar results using the ancestral allele and context from the 1000G reconstruction of the ancestral human genome sequence (41) (SI Appendix, Fig. S2). We include multiallelic sites (∼6% of the data) by counting the multiple derived alleles separately as if they had occurred at separate bi-allelic sites with the same major allele (SI Appendix, Fig. S3). To obtain the diversity level for each mutation type within a genomic compartment, we divide the number of segregating sites of a particular type by the number of mutational opportunities, i.e., sites where a single change could have given rise to that mutation type; this approach accounts for base composition within a compartment.

To compare mutation types across two genomic compartments, we normalize the diversity for each mutation type by the total diversity within each compartment. In this way, we control for the effect of population genetic processes that affect diversity across compartments but do so evenly across all mutation types, and isolate differences in the mutation spectrum; this step is particularly important for comparisons between the X chromosome and autosomes. For each of 96 mutation types, we test if the observed relative diversity in the two compartments differs from what would be expected by chance. To this end, we designate one of the two compartments as the “test” and the other as the “reference” compartment. Our null expectation is that the number of mutations of a particular type in the test compartment is binomially distributed with a mean value proportional to the observed diversity for that type in the reference compartment, adjusted for overall differences in diversity between the two compartments (SI Appendix). Mutation types are considered significantly different in their frequencies between the two compartments if the two-tailed p-value from the binomial test is below the Bonferroni-corrected 5% significance threshold. This approach implicitly ignores sampling error in the estimate of diversity of the designated reference compartment; we verify that our results are insensitive to this assumption by using alternative approaches to calculate significance that do not make this assumption, but have other limitations (SI Appendix; Dataset S1). We consider the effects of highly mutable types on the distribution of other mutation types (SI Appendix; Fig. S4); we also consider possible differences in sequencing error rates between compartments, and replicate our findings in two alternate datasets (SI Appendix; Fig. S5-S7).

## Results and Discussion

Biochemical properties vary along the genome, both on autosomes and the X chromosome. In turn, sex-specific influences from the germline are the same across autosomes, but differ between the X chromosome and autosomes. We therefore first compare autosomal compartments with distinct biochemical features to illuminate biochemical influences on the mutation spectrum. Then, by comparing compartments across the X chromosome and autosomes and accounting for average biochemical differences between them, we disentangle sex-specific and biochemical influences on the mutation spectrum.

### Replication timing and its covariates influence the germline mutation spectrum

We consider autosomal compartments that differ with regard to specific biochemical properties in the germline. In cases where these data are unavailable for germline tissue and we are limited to somatic cell lines, we focus on biochemical features that have broadly similar distributions across tissue types. Replication timing is consistently an important predictor of local mutation rates (Stamatoyannopoulos et al. 2009; Smith, Arndt, and Eyre-Walker 2018; Chen et al. 2017) in both the soma and the germline, and broad-scale replication timing maps are relatively concordant across tissues (43, 44) (Fig. S8). The observed mutagenic effect of late replication has been hypothesized to be due to a decline in the efficacy of mismatch repair with delayed replication, less time for repair, or the accumulation of damage-prone single-stranded DNA at stalled replication forks (26, 42).

To assess if replication timing affects the germline mutation spectrum, we compare autosomal regions that differ in their replication timing using available data from LCL and H9-ESC cell lines (43, 45). As expected, almost all mutation types are significantly enriched in late replicating regions relative to early replicating regions (Fig. 2a, Fig. S9a). In particular, we observe a substantial enrichment of C>A and T>A mutations in late replicating regions, a pattern also observed by Carlson et al., 2018 in a different sample of rare variants. Moreover, the mean replication timing in 1 Mb windows across the genome explains ∼60% of the variation in C>A and T>A enrichment in those windows relative to the autosomal average and between 2% and 26% for all other mutation types (p << 10^-5^), suggesting that these two mutation types are particularly sensitive to replication timing (Fig. 2b, Fig. S9b, SI Appendix).

**Fig. 2.**
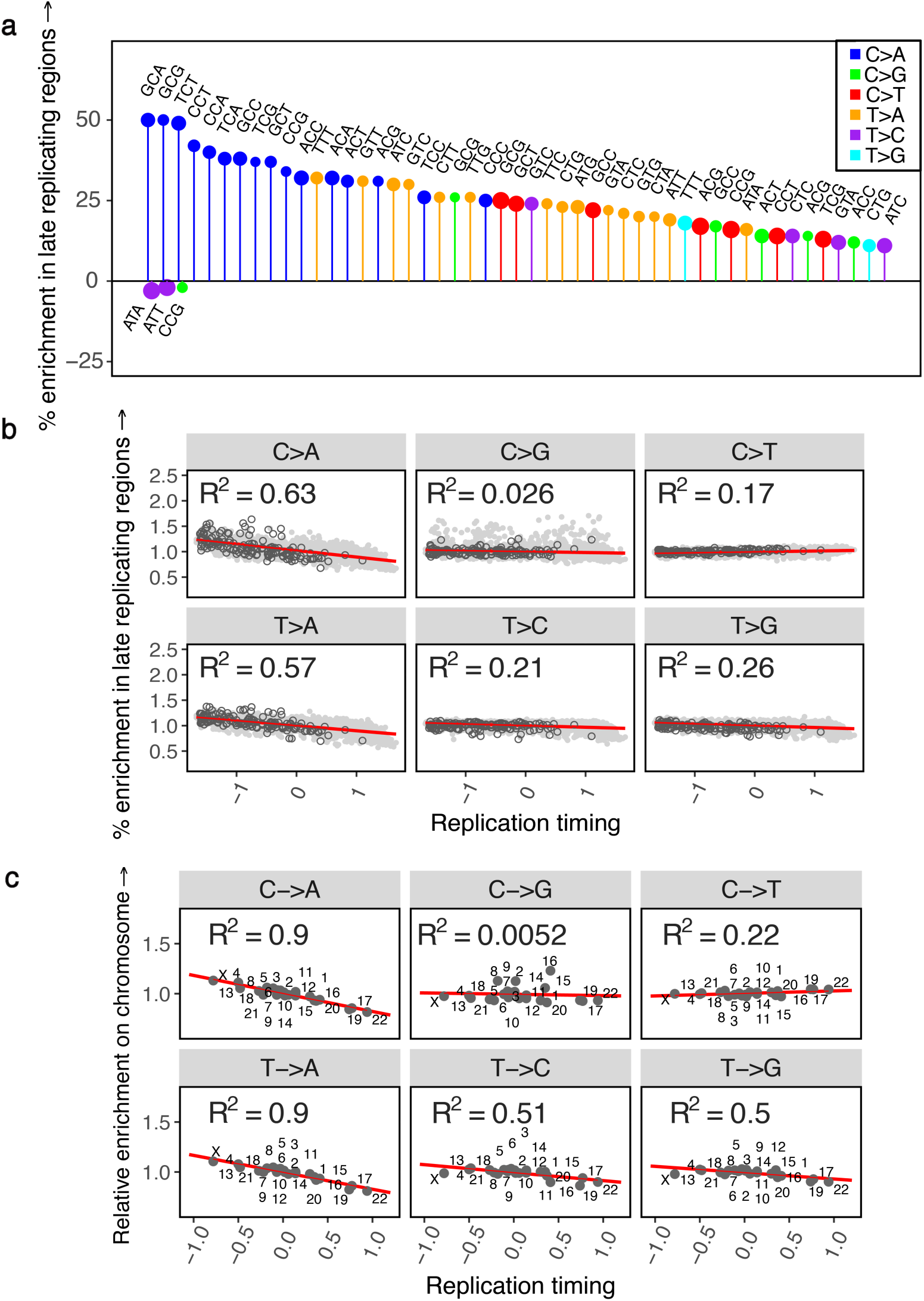
The effect of replication timing on the mutation spectrum at different scales, using replication timing scores from the LCL cell line. (a) Comparison of the spectrum of 96 mutation types in late replicating (score <= −0.5) autosomal regions relative to early replicating (score >= 0.5) autosomal regions. Positive and negative effects have been separately ordered by effect size from left to right; only the top 50 significant positive and negative effects are shown for legibility. The size of the circle reflects the number of mutations of that type. (b) For each of six mutational classes, the enrichment in 1Mb windows relative to all other autosomal windows combined, ordered by the mean replication timing. Positive replication timing scores indicate earlier than average replication. Autosomal windows are shown in solid light grey circles; windows on the X chromosome have been overlaid in black hollow circles. (c) For each of six mutational classes, enrichment on individual autosomes and X relative to all other autosomes combined, ordered by the mean replication timing. Positive replication timing scores indicate earlier than average replication.

Because replication timing is correlated with multiple genomic features, including higher order chromatin structure, epigenetic modifications, and in particular, DNA methylation at CpG sites, some of the observed patterns could be reflective of these processes rather than replication per se. To assess the marginal impact of CpG methylation on the effect of replication timing, we consider early and late replicating regions within and outside CpG islands, which are regions of CpG hypomethylation across tissue types (46, 47). We find that at both CpG sites inside and outside islands, C>A mutations are enriched in late replicating regions (Fig. S10), suggesting that this signal is not due to differences in methylation. Moreover, we also observe this pattern at non-CpG sites (Fig. S10).

The association of C>A and T>A mutations with replication timing does not necessarily imply that they are “replicative” in origin, i.e., due to errors directly introduced by the replication machinery while copying intact DNA, as they could also reflect greater unrepaired damage in later replicating regions (26). In particular, since C>A mutations are a known consequence of oxidative damage in somatic tissues (Alexandrov et al. 2018; David, O’Shea, and Kundu 2007; Neeley and Essigmann 2006; De Iuliis et al. 2009; Alexandrov et al. 2013), it is plausible that these mutations accumulate in regions of late replication due to greater damage to exposed single-stranded DNA, or poorer repair in these regions.

Considering other factors shown to influence mutation patterns, we recover a known signature of CpG methylation: transitions at CpG sites (C>T mutations in the ACG, CCG, GCG, and TCG trinucleotide contexts), which are thought to be due to the spontaneous deamination of methyl-cytosine to thymidine, are highly depleted in the hypomethylated CpG islands compared to the rest of the genome (Fig. S11a). Similarly, we detect an increase in C>G mutations in a subset of autosomal regions previously shown to be enriched for clustered C>G de novo mutations (Fig. S11b). This C>G signature is thought to reflect inaccurate repair of spontaneous damage-induced double-strand breaks in the germline (29, 51).

Importantly, the impact of these biochemical features on mutation does not average out across chromosomes. Comparing individual autosomes to all other autosomes reveals ubiquitous variation in the mutation spectrum at the chromosome-level (Fig. S11c). In particular, individual chromosomes that replicate later on average show greater enrichment of C>A and T>A mutation types: differences in mean replication timing for individual autosomes explain ∼90% of the variation in C>A and T>A enrichment at the chromosome level (p << 10^-5^), while they explain ∼50% or less for other mutation types (Fig. 2c). These results demonstrate that replication timing, and potentially other genomic features such as methylation and propensity for accidental double strand break damage, lead to chromosome-level differences in diversity, hinting at some plausible sources for observed but unexplained chromosome-level differences in average divergence (37).

### Sex-specific influences on the mutation spectrum are subtle but likely ubiquitous

Next, we assess the impact of sex on the germline mutation spectrum by comparing mutational patterns on the X chromosome and autosomes. The X chromosome is disproportionately exposed to mutational processes in the female germline; viewed from a population perspective, there are more X chromosomes in females than in males, but the same number of autosomes in both. Thus, mutation types that arise more commonly in the female germline are expected to be enriched (and mutation types that arise more commonly in the male germline depleted) on the X chromosome relative to autosomes. We account for population-level properties that may affect the mutation spectrum differently on the X and autosomes (SI Appendix). Having done so, we find most mutation types to be differentially enriched on the X and autosomes (Fig. 3a).

**Fig. 3.**
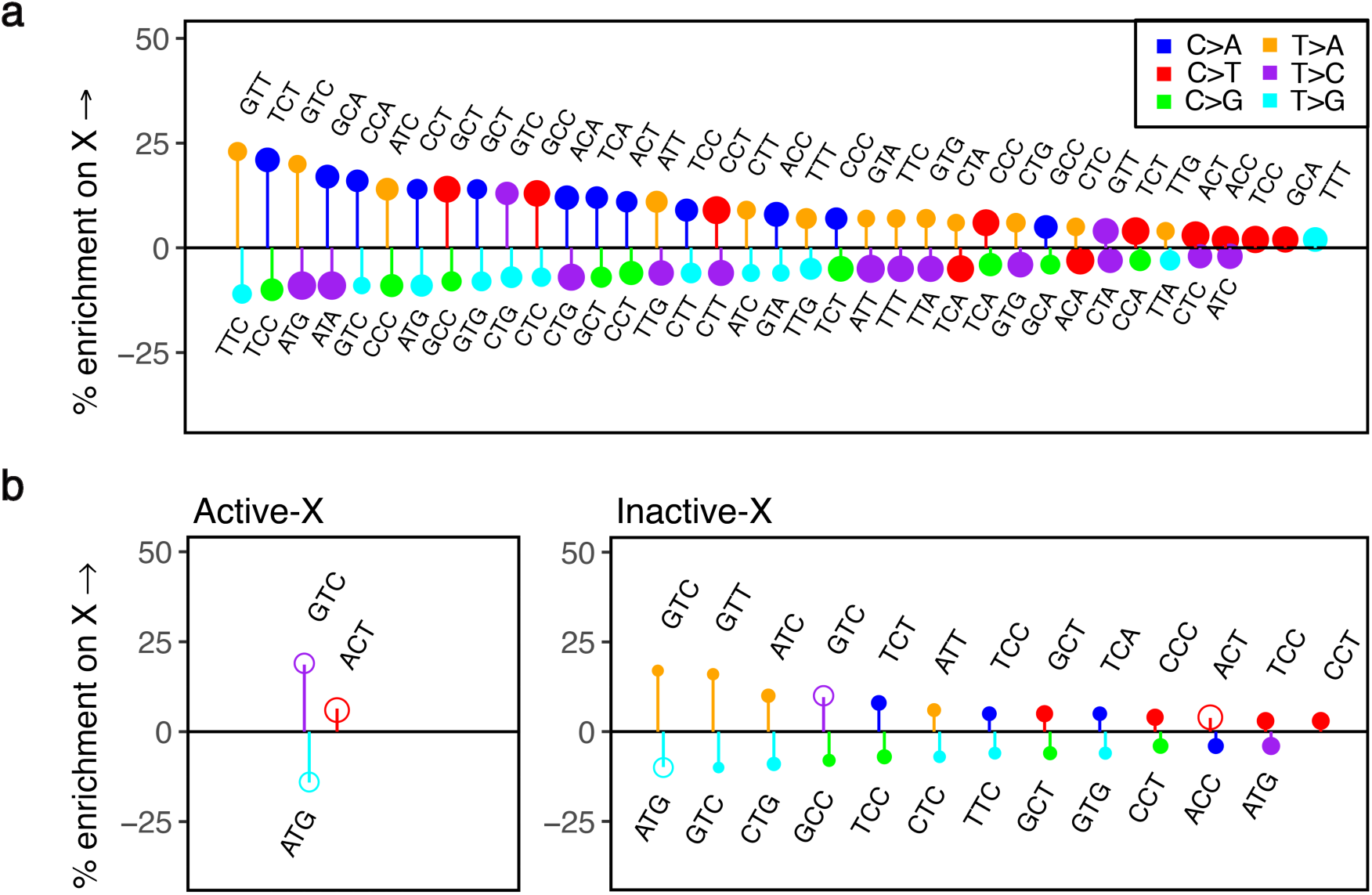
Comparison of the mutation spectrum on the X chromosome and autosomes. The pseudoautosomal region (PAR) and CpG sites are excluded from this analysis. Only significant differences are shown. Positive and negative effects have been separately ordered by effect size from left to right. The size of the circle reflects the number of mutations of that type. (a) Enrichment of mutation types on the X chromosome relative to autosomes. (b) Enrichment of mutation types in the genic compartment of the X chromosome that escapes inactivation, relative to genic regions on autosomes. Hollow circles represent mutation types enriched (or depleted) in both the escaped (active) and inactive compartments of the X relative to autosomes. (c) Enrichment of mutation types in the genic X chromosome compartment that undergoes X-inactivation, relative to genic regions on autosomes. Hollow circles represent mutation types enriched (or depleted) in both the escaped (active) and inactive compartments of the X relative to autosomes. The larger number of significant differences in (c) compared to (b) likely reflects at least in part the approximately five-fold greater amount of data in the inactive versus the active genic regions of the X chromosome.

Importantly, however, these X-autosome differences do not only reflect differences in male and female mutational processes; given the substantial effect of biochemical features on mutational patterns observed at the chromosomal scale, they also potentially reflect differences in the distribution of these biochemical features on the X chromosome and autosomes. For instance, in de novo mutation studies (29, 32), C>A mutations are found to arise more often in males, suggesting that they should be depleted on the X. Instead, they are found enriched on the X chromosome relative to autosomes (Fig. 3a). A possible explanation is that the X accrues excess C>A mutations because it replicates late in the germline. C>A mutations are known to be associated with oxidative damage (Alexandrov et al. 2018; David, O’Shea, and Kundu 2007; Neeley and Essigmann 2006; De Iuliis et al. 2009; Alexandrov et al. 2013), which remains unrepaired in sperm (Smith et al. 2013), and is likely repaired at or before the first cell division in the zygote (Harland et al., 2017; Huang et al., 2014; Ju et al., 2017). Late replication of the X chromosome at this stage, perhaps due to the inactive status of the paternally inherited X in female embryos (52), could then indeed be expected to result in an enrichment of C>A mutations on the X relative to autosomes, despite a primarily male source of damage. This example underscores that accounting for the X-specific effects of biochemical features is key to uncovering true sex differences in X-autosome comparisons.

One well-characterized idiosyncratic property of the X is X-inactivation, which is associated with X-specific changes in methylation, transcriptional activity, and notably, replication timing: because the inactive X chromosome exhibits a significant lag in replication, on average the X replicates later than autosomes (53). Though X-inactivation is a short-lived process in the germline—limited to early embryogenesis in females, and brief meiotic and post-meiotic periods in males (54–57)—it could nevertheless lead to observable differences in the mutation spectrum between different regions of the X. The “active” compartment of the X chromosome, i.e., the approximately 15% of the transcribed X that constitutively escapes inactivation across tissues (58, 59) may therefore differ in its mutation spectrum from the rest of the X. Comparing autosomes with the inactive and active regions of the X, we find T>C mutations at GTC sites and C>T types at ACT sites enriched in both active and inactive regions of the X relative to autosomes and T>G mutations at ATG sites depleted both in the active and inactive regions of the X relative to autosomes (Fig. 3b and 3c, Fig. S12). Since these cases cannot be attributed to X-inactivation and are enriched (or depleted) concordantly on compartments of the X chromosome that differ in their replication timing, methylation levels and other features, they are strong candidates for true sex differences in mutation. Given that the genic compartment known to escape inactivation across tissues is a small fraction of the X chromosome, there are likely many more subtle ones that we miss.

A complementary approach to minimizing X-specific biochemical influences on the mutation spectrum of the X in X-autosome comparisons is to consider regions of the X chromosome that are comparable to autosomes in their average replication timing. The replication timing on the X chromosome across multiple human cell lines depends on whether one of the X chromosomes is inactivated (Fig. S8; SI Appendix) (Allegrucci and Young 2007; Vallot et al. 2015; Patel et al. 2017; Ryba et al. 2010; Hiratani et al. 2010; Tang et al. 2016). This observation suggests that controlling for replication timing differences between the X chromosome and autosomes may also control for the effects of other correlated features, including those associated with X-inactivation. Using this approach, all three mutation types that we highlight as putative sex differences based on their differential enrichment in the active compartment of the X relative to autosomes are also observed as significant differences between the X chromosome and autosomes (Fig. 4a, Fig. S13). That we find the same types with this complementary approach provides further evidence that they are true sex differences.

**Fig. 4.**
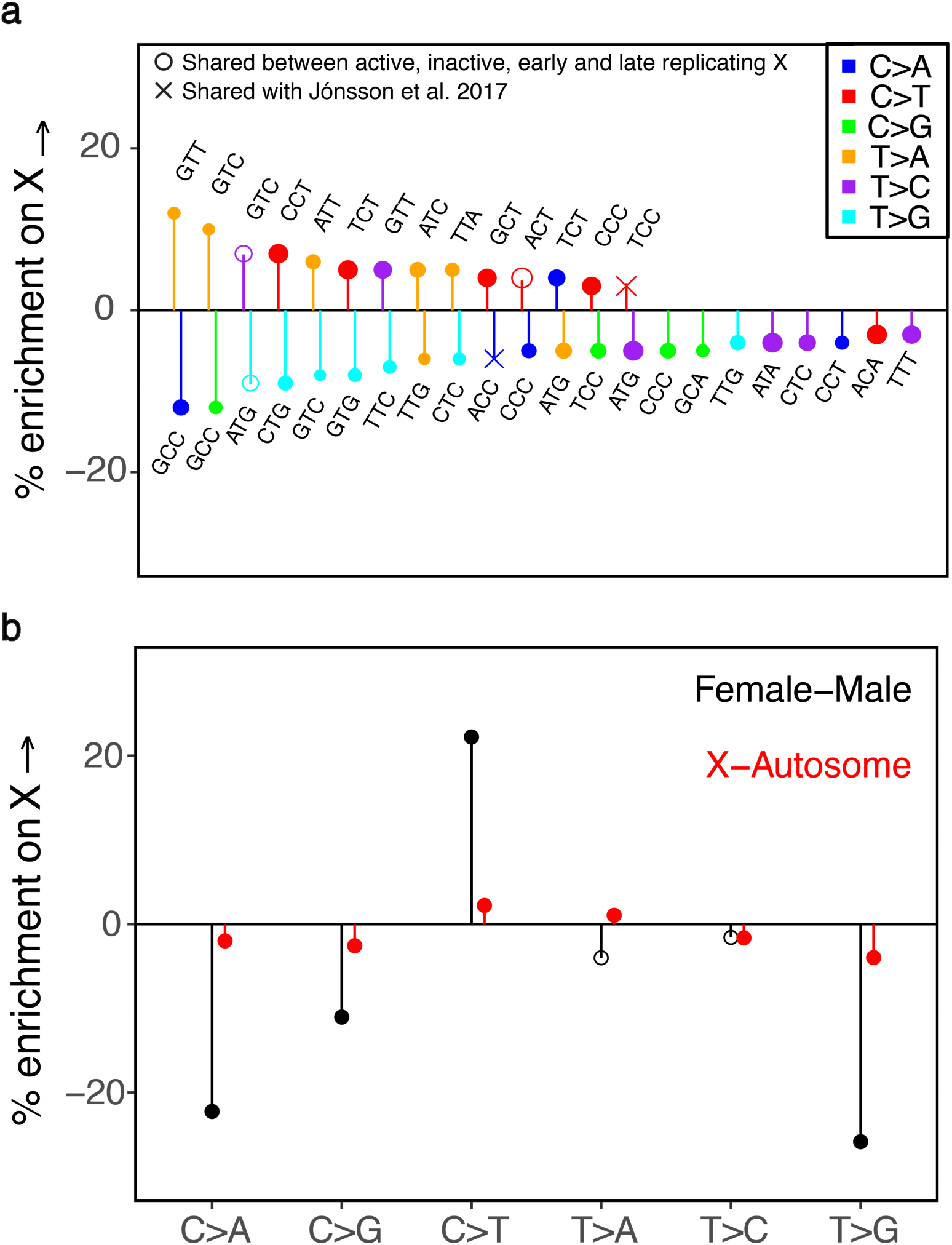
The mutation spectrum on the X and autosomes matched for average replication timing. The pseudoautosomal region (PAR) and CpG sites are excluded from this analysis. (a) Comparison of the mutation spectrum on the X chromosome and autosomes matched for average replication timing. Only significant differences are shown. Positive and negative effects have been separately ordered by effect size from left to right. Hollow circles represent mutation types also enriched (or depleted) in both the active and inactive compartments of the X relative to autosomes. Crosses denote mutation types reported as significant sex differences by Jónsson et al., 2017. (b) The X-autosome spectrum for six mutation classes, controlling for mean replication timing (in red), compared to known male female differences from Jónsson et al., 2017 (in black). Solid points are statistically significant differences at the 5% level, accounting for multiple tests.

We also detect a number of additional differentially enriched types between X and autosomes after controlling for replication timing differences (Fig. 4a); many of these types are enriched concordantly in early and late replicating regions of the X relative to autosomes (Fig. S14). Assuming that a majority of X-specific effects are accounted for when we control for replication timing, these types can also be considered putative sex differences. In that respect, it is noteworthy that C>A mutations are enriched in inactive or late replicating regions of the X, but depleted in the active or early replicating regions of the X, when compared to autosomes (Figs. S12, S14). This pattern is what we would expect from the combined influences of a male bias and an effect of replication timing on C>A mutations, as we suggested earlier.

We further assess these putative sex-specific signatures by comparing them to results from the largest human pedigree study of de novo mutations to date (29). Among the six broad mutational classes, Jónsson et al. find C>T mutations significantly enriched in maternal, and C>A, C>G, and T>G mutations relatively enriched in paternal de novo mutations. The mutational patterns we observe on the X chromosome and autosomes after controlling for differences in replication timing are consistent with these effects: we find C>T mutations significantly enriched and C>A, C>G, and T>G classes significantly depleted on the X chromosome relative to autosomes (Fig. 4b; Fig. S13). Jónsson et al. also find three mutation types in their trinucleotide context (TCC>TTC, ACC>AAC, ATT>AGT) as significant sex differences: of these we find two as significant X-autosome differences. As expected, the maternally enriched TCC>TTC type is relatively enriched on the X chromosome, and the paternally enriched ACC>AAC type is relatively enriched on autosomes (Fig. 4a). We do not observe the third type as differentially enriched on the X and autosomes, possibly because there are genomic features specific to the X that mask its enrichment in females.

In turn, the types that we identify as putative sex differences from the comparison of X active, X inactive and autosomes are not reported as significant sex differences in Jónsson et al. (2017). The reason may be that most of them reflect subtle X-autosome differences, with X-enrichment or depletion in the range of 5-10%. Translating these enrichments into a difference between males and females requires a full population genetic model, including assumptions about demography and life history (63). Nonetheless, such subtle X-autosome differences likely correspond to small sex differences that current de novo studies are underpowered to detect.

### Accidental and a subset of meiotic double-strand breaks in the germline have the same mutagenic impact

In the preceding section, we suggested a plausible mechanism through which local biochemical influences and sex-specific properties of the germline jointly influence the distribution of C>A mutations on the X chromosome and autosomes. Here we highlight another mutation type, C>G, which is also distributed in a sex-specific and chromosome-specific manner, but is largely insensitive to replication timing.

As recently reported, clustered C>G de novo mutations are concentrated in particular autosomal regions, and the number of such mutations transmitted in each generation increases exponentially with maternal age at reproduction (29, 51). Maternal age at reproduction determines the duration of oocyte arrest, since females are born with their entire complement of oocytes, which remain in dictyate arrest until ovulation. Based on the sex-specific patterns of accumulation with age and genomic properties of these mutations, the authors speculated that the C>G clusters could be due to the more frequent spontaneous occurrence of damage-induced double strand breaks (DSBs) in some genomic regions and an increasing rate of such damage in older oocytes. In this view, C>G mutations are associated with accidental double-strand break damage in both males and females.

Accidental damage is not the only source of double strand breaks in the germline, however: during meiosis, double strand breaks are deliberately induced along the genome, through targeting of PRDM9-binding motifs (64, 65). These DSBs are repaired through the homologous recombination pathway: a small minority are resolved through crossovers (COs), which involve exchanges of large segments between homologous chromosomes, and the rest are thought to be repaired through non-crossover gene conversion events (NCOGCs), though another small subset may involve non-homologous end joining and other mechanisms (66, 67). Potentially, these meiotic DSBs could have a mutagenic impact similar to that of spontaneous double-strand breaks; however, a clear mutational pattern common to both has not been seen to date. For instance, using DMC1 ChIP-Seq data from human spermatocytes, Pratto et al., 2014 observed C>G enrichment to a small degree around male autosomal hotspots (68), but the source of these types was not discussed further by the authors, and is potentially due to overlap with regions of clustered de novo C>G mutations reported by Jónsson et al., 2017. Another recent study did not find de novo C>G mutations enriched within autosomal crossover hotspots identified in pedigree studies (69). We test if there is indeed an enrichment of C>G mutations associated with meiotic DSBs by comparing the mutation spectrum within and outside hotspots on autosomes; we use DMC1 hotspots in males and crossover hotspots in females because we do not have a map of DMC1-binding in female gametes. Our results are consistent with previous observations: we do not observe C>G enrichment in autosomal hotspots for males or females once we exclude regions of clustered de novo C>G mutations (Figs. S11b, S15a, S15b).

We next consider the X chromosome, which in females recombines like an autosome, but in males is in the unusual position of having no homolog outside the pseudoautosomal region (PAR). In males as in females, meiotic DSBs are nonetheless made both inside and outside the PAR (68, 70–72). Properties of recombination events on the X and autosomes differ markedly between sexes however: notably, the pseudoautosomal region 1 (PAR1), a 2.6 Mb region on the X chromosome, experiences an obligate crossover in males, but normal levels of recombination in females (70, 73–75) and in male germ cells, DSBs are repaired late on the X chromosome relative to autosomes (68, 70–73). These considerations raise the possibility that mutational patterns in hotspots on the X chromosome in males may reflect these sex-specific features of recombination and behave differently relative to autosomal hotspots in males, and relative to both X and autosomes in females. To explore this hypothesis, we compare mutation patterns on autosomes to those in PAR1, which is exposed to the male and female germlines to the same degree as autosomes (since two copies are carried by both males and females), and does not undergo X-inactivation (76). We find that C>G mutation types are systematically enriched on the PAR1 relative to autosomes (Fig. 5a), indicating that repair of meiotic double-strand breaks in this region in males is associated with C>G enrichment.

**Fig. 5.**
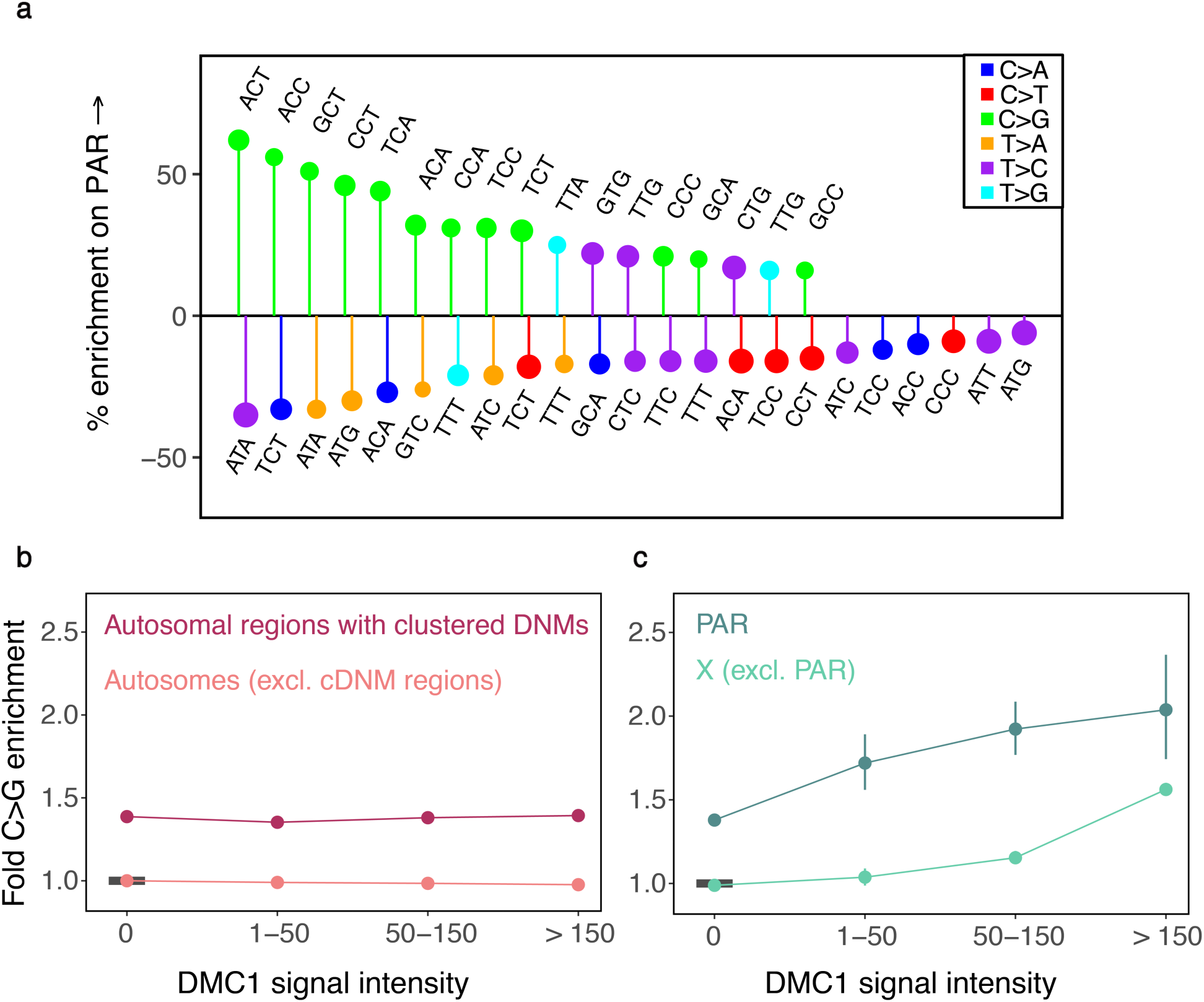
Distribution of C>G mutations in genomic compartments relative to autosomes. CpG sites are excluded from these analyses. (a) The mutation spectrum on PAR1 relative to autosomes. Only significant differences are shown. Positive and negative effects have been separately ordered by effect size from left to right. (b) Enrichment of C>G mutations in DMC1 hotspots of varying intensity on autosomes. For each estimate, the 95% confidence interval from a binomial test is represented by the vertical bars (and is sometimes too small to be apparent). The horizontal black bar shows the reference, namely autosomes outside DMC1 hotspots and excluding regions rich in clustered de novo mutations (identified by Jónsson et al., 2017). (c) Enrichment of C>G mutations in DMC1 hotspots of varying intensity on the X chromosome, relative to autosomes outside DMC1 hotspots and excluding autosomal regions rich in clustered de novo mutations. For each estimate, the 95% confidence interval from a binomial test is represented by the vertical bars. The reference, same as for (b), is denoted by a horizontal black bar.

We further characterize the source of the C>G enrichment using DMC1 ChIP-Seq data from human spermatocytes (68). The DMC1 signal reflects intermediates in the homologous recombination pathway; high levels of DMC1-binding can reflect either an increased frequency of double strand breaks (hotspots of greater intensity) or a greater duration of intermediates, i.e., a longer time to repair (68, 70). Using these data, we find that there is clear C>G enrichment not only in PAR1, but also in hotspots on the X chromosome outside PAR1; moreover, the enrichment increases with the strength of the DMC1 signal (Fig. 5b-5c, Fig. S15c). That we observe C>G enrichment in male hotspots on the X chromosome but not in male hotspots of similar average intensities on autosomes (Fig. 5b-5c, Fig. S15a) or in female crossover hotspots on autosomes (Fig. S15b), leads us to speculate that the predominant source of this C>G signature is the delay in repair of DSBs on the X chromosome relative to autosomes in male meiosis. We note that because hotspots detected in spermatocytes could also be used in female meiosis (77), without a female-specific map of DMC1 binding, we cannot exclude the possibility that C>G enrichment is also associated with recombination on the X in females; however, the observed C>G enrichment in the strongly male-biased hotspot PAR1 (Fig. 5) supports our conjecture of a male-specific impact of meiotic DSBs on the X, at least in this region.

One possibility is that the enrichment of C>G mutations stems from a switch in the repair machinery late in meiosis (78–80); DSBs still not repaired by this stage may be repaired by a more mitotic-like repair pathway, which could potentially be more mutagenic (80, 81). Notably, the source of the C>G signature found by Jónsson et al., 2017 in specific autosomal regions could also be late repair; indeed, if these areas reflect damage, as the authors surmise, they may only undergo repair later in meiosis. Thus, a shared biochemical pathway may underlie the mutagenic impact of both spontaneous and a subset of meiotic DSBs. Moreover, our results illustrate a subtle sex-specific mutagenic effect of meiotic recombination, whereby the repair of meiotic DSBs on the X specifically in males gives rise to C>G mutations; in that sense, components of the recombination machinery that are involved in late repair of double-strand breaks are sex-specific modifiers of mutation.

### Implications

By comparing the mutation spectrum across different compartments of the genome, we identify putative signatures of sex differences in the germline and plausible biochemical sources of mutagenesis. Notably, we show that replication timing affects the mutation spectrum along the genome and find a mutagenic effect of meiotic recombination that is both sex-specific and X-specific, revealing an appreciable effect of double-strand breaks, both accidental and deliberate, on the mutation spectrum.

Interestingly, our analysis suggests that signatures of sex differences in the germline are likely abundant, but their contributions to the mutation spectrum are subtle relative to those of biochemical processes shared in the two sexes. This finding is hard to reconcile with the idea that male mutations are mostly replication-driven whereas female mutations reflect a large contribution of spontaneous damage, as then we might expect substantially different types of mutations inherited from mothers and fathers. Instead, consistent with a greater role of spontaneous damage and its repair in both male and female germlines (51), our results are most readily explained if male and female mutational mechanisms are overall highly similar, underpinned by the shared mechanisms associated with replication, transcription, methylation, and recombination, and other sources of damage. Subtle differences in the mutation spectrum between males and females could then be expected to arise due to sex-specific rates of damage and repair at different stages in germline development.

These sex differences in germline mutation are modulated by life history traits of males and females. As one example, the proportion of C>G mutations transmitted in a single generation increases with the age of the mother (29, 51). Indeed, even when there are no sex-differences in the biochemical process itself, much of the biochemical machinery that influences mutation must in theory have subtle sex-specific effects, simply because sex-specific life history traits modulate exposure to biochemical influences differently in males and females. Changes in life history traits, or in the frequency of variants associated with sex-specific life history traits over evolutionary time could then change the proportion of particular mutation types and thus alter the mutation spectrum over time. Together with other sex-specific modifiers of mutation, life history traits likely play a role in the evolution of the mutation spectrum not only on autosomes, but also on the X chromosome relative to autosomes.

In this respect, we note that a number of recent studies have shown that the mutation spectrum changes slightly across populations (3, 82–84). These findings have largely been attributed to biochemical modifiers of mutation that alter the relative rates of different mutation types by influencing the biochemical process of error/repair over time. Our results highlight that life history traits and other sex-specific modifiers could potentially result in the same kinds of changes in the mutation spectrum and the mutation rate over time. Moreover, parental ages of reproduction explain a large proportion of the observed mutation rate variation among ∼1500 individuals at present (29). Variants that contribute to sex-specific life history (85, 86) may therefore be a useful starting point to identify genetic sources of inter-individual variation in the mutation rate in humans.

Beyond these insights into mutagenesis, our analysis makes clear that X-autosome comparisons of mutation patterns cannot be taken as directly reflective of germline sex differences. Though historically comparisons of the sex chromosomes to autosomes have been taken to reflect only the effects of sex, mutation patterns on the X chromosome in fact reflect a convolution of X chromosome specific effects and sex. Taking this point into consideration may help to explain, for instance, why estimates of the male bias in mutation for CpG sites from phylogenetic studies that used X-autosome comparisons are much lower (87) than those obtained directly from male-female differences in de novo mutation data (12, 29).

## Acknowledgements

We thank Guy Amster, Ziyue Gao, Priya Moorjani, Itsik Pe’er, Jonathan Pritchard, Guy Sella, Arbel Harpak, Felix Wu, and additional members of the Przeworski lab for helpful discussions and/or comments on a draft version of the manuscript and Priya Moorjani, Konrad Karczewski, and Kelley Harris for assistance with gnomAD and SGDP data sets. We are also grateful to comments from three anonymous reviewers on an earlier draft. This work was supported by R01 GM122975 to M.P.

