## Supplementary Methods and Figures for "Signatures of replication timing, recombination and sex in the spectrum of rare variants on the human X chromosome and autosomes"

**This PDF file includes:**

Supplementary Methods
Figs. S1 to S15

**Other supplementary materials for this manuscript include the following:**

Dataset S1 Supplementary_Methods_Tables_1-3.xlsx
Dataset S2 Supplementary_Data_for_Figures.xlsx

**Supplementary Methods**

### Data

The data used and their sources are summarized in the tables below:

| **Variants** | |
|---|---|
| gnomAD | gnomAD release 2.0.1 (http://gnomad.broadinstitute.org) |
| SGDP | SGDP-Lite (http://reichdata.hms.harvard.edu/pub/datasets/sgdp/) |
| UK10K | ALSPAC and TwinsUK (https://www.uk10k.org/data_access.html) |

| **Genomic features** | |
|---|---|
| Replication timing | LCL (http://mccarrolllab.org/wp-content/uploads/2015/03/Koren-et-al-Table-S2.zip).<br>H1- hESC (wgEncodeFsuRepliChipH1hescWaveSignalRep1);<br>H7-hESC (wgEncodeFsuRepliChipH7esWaveSignalRep1);<br>H9-hESC (wgEncodeFsuRepliChipH9esWaveSignalRep1) from https://genome.ucsc.edu/encode/ |
| CpG islands | http://www.haowulab.org/software/makeCGI/model-based-cpg-islands-hg19.txt |
| DMC1 ChIP-seq | Human spermatocytes (GEO Accession GSE59836) from http://www.ncbi.nlm.nih.gov/geo/ |
| Recombination rate | Standardized female recombination maps (https://www.decode.com/addendum/) |
| Clustered DNM regions | Supplementary Table 12 (https://www.nature.com/articles/nature24018) |
| Genic regions | Gencode v19 genes (ftp://ftp.ebi.ac.uk/pub/databases/gencode/Gencode_human/release_19/gencode.v19.annotation.gtf.gz) |
| Genic regions of X-inactivation and escape | Supplementary Table 13 (https://www.nature.com/articles/nature24265) |
| Pseudoautosomal region (PAR) | http://genome.ucsc.edu/cgi-bin/hgGateway |

| **Reference genome** | |
|---|---|
| hg19 | https://www.ncbi.nlm.nih.gov/assembly/GCF_000001405.13 |

**Delineating the set of variants in the gnomAD dataset.**

We used publicly available whole genome SNP data from 15,496 individuals compiled and made available by the Genome Aggregation Database (gnomAD), which includes 9,256 Europeans and 4,368 African or African-American individuals (1). We restricted our dataset to a set of good quality SNPs that passed the baseline quality filter provided by gnomAD, such that there was at least one individual at each site with a high-quality non-reference genotype: quality-adjusted allele count (AC) > 0, or equivalently, Filter = "Pass", DP > 10, GQ > 20, and AB > 0.2 for heterozygotes. We excluded sites that overlap with indels and CNVs. We retained multi-allelic sites (6.5% and 5% of the data on the autosomes and X respectively).

Since our goal was to compare genomic compartments, including those on the X chromosome, we matched the number of X chromosomes and autosomes in our sample by limiting our analysis to the 6,930 female individuals in the sample (using the quality-adjusted female allele counts provided). Additionally, we imposed separate filters on the X chromosome and autosomes so that only variants with an allelic depth within one standard deviation of the mean allelic depth on the X chromosome (13760 +/- 512) and autosomes (13753 +/- 562), respectively, were retained in the sample (only about 2.8% of data from both the X-chromosome and autosomes is lost at this step).

**Calculating diversity levels for 96 mutation types.**

Most variants in the gnomAD dataset are extremely rare: about 64% are singletons and doubletons (i.e., variants seen once or twice in the sample). Only ~10% of variants are at frequency 1% or greater (Fig. 1b); their inclusion does not affect our qualitative results, since they are a small subset of the data, and their mutation patterns are largely the same as variants at lower frequencies (Supplementary Figure 1); we therefore retain them.

The variants in the gnomAD dataset are called with respect to the human reference (hg19). We instead polarized to the major allele in the full sample of 15,496 individuals, so that the minor allele was treated as derived. At multi-allelic sites, we counted the multiple derived alleles separately as if they had occurred at separate bi-allelic sites with the same major allele. We obtained similar results (Supplementary Figure 2) using the ancestral allele and context from the 1000G reconstruction of the ancestral human genome sequence (2).

As is standard (e.g., Alexandrov et al. 2013; Harris 2015; Harris and Pritchard 2017), we extracted the single base pair flanking sequence on each side of the variant position using the portion of the hg19 reference callable in gnomAD to obtain mutations in their trinucleotide context, and combined mutations in reverse complement classes (for example, the ACG>ATG and CGT>CAT classes were collapsed into the former) to obtain 96 mutation types. To obtain diversity levels for each of the 96 mutation types, we divided the number of segregating sites of a particular type by the number of mutational opportunities at that type of site, where mutational opportunities are defined as sites at which a single change could have given rise to the mutation type under consideration (note that there are three mutational opportunities at each base pair in the genome). By dividing the number of mutations by the number of possible mutations in each genomic compartment, we account for base composition differences between compartments.

**Comparing diversity levels between genomic compartments**

In comparing a pair of genomic compartments, we took differences in their population genetic properties into account. By normalizing the diversity levels for each mutation type by overall diversity levels for the two compartments, we controlled for the effect of population genetic processes that affect diversity across compartments but are expected to do so evenly across all mutation types, allowing us to isolate differences in the mutation spectrum. This normalization is particularly relevant for comparisons between the X chromosome and autosomes as, for the same sample size, there are more neutral segregating sites expected on autosomes: because the autosomes spend more time in the male germline relative to the X chromosome, they have a higher overall mutation rate, as well as a slightly larger effective population size due to differences in the genealogical process between the X-chromosome and autosomes.

Suppose that in an arbitrary genomic compartment "a" with $n$ total mutations ("segregating sites") of all types and $n_i$ mutations of type $i$, the proportion of mutation type $i$ is $n_i/n$. If there are $d_i$ potential sites ("mutational opportunities") at which this mutation type could occur, out of $d$ total sites in the compartment, the normalized proportion (or normalized diversity) of mutation type $i$ is:

$$r_{i(a)} = \frac{(n_{i(a)}/d_{i(a)})}{(n_a/d_a)} = \frac{\pi_{i(a)}}{\pi_{(a)}}; \qquad \mathrm{n}_{(a)} = \sum_i n_{i(a)}, \mathrm{d}_{(a)} = \sum_i d_{i(a)}$$

The relative enrichment of this mutation type in compartment a, compared to another compartment b, is then:

$$R_{i(ab)} = \frac{r_{i(a)}}{r_{i(b)}} = \frac{\pi_{i(a)}/\pi_{(a)}}{\pi_{i(b)}/\pi_{(b)}} = \frac{(n_{i(a)}/d_{i(a)})/(n_{i(b)}/d_{i(b)})}{(n_a/d_a)/(n_b/d_b)}$$

In particular, when the two compartments under consideration are the X chromosome and autosomes, normalizing by overall diversity allows us to take into account the different population level effects of demography and life history on these compartments, captured in the effective population size ($N_e$) parameter (under some simplifying assumptions, e.g., no multiple hits), and isolate differences in the mutation spectrum:

$$\frac{\bar{\pi}_{i(X)}/\bar{\pi}_{(X)}}{\bar{\pi}_{i(A)}/\bar{\pi}_{(A)}} = \frac{\mu_{i(X)} \cdot N_{e(X)}/\mu_{(X)} \cdot N_{e(X)}}{\mu_{i(A)} \cdot N_{e(A)}/\mu_{(A)} \cdot N_{e(A)}} = \frac{\mu_{i(X)}/\mu_{(X)}}{\mu_{i(A)}/\mu_{(A)}}$$

where $\bar{\pi}$ is mean diversity and $\mu$ denotes the mutation rate.

In large samples with recurrent mutations (i.e., repeat mutations or "multiple hits" at the same site), normalizing by overall diversity does not account fully for population level effects, particularly for sites with high mutation rates. In particular, at the highly mutable CpG sites, recurrent mutations are frequently expected in a sample of this size (1, 6). Because autosomes have a slightly larger effective population size and higher mutation rate compared to the X, we expect more recurrent mutations at these sites on autosomes. Although we include multi-allelic sites in our analysis and can therefore count mutations to three different alleles at a site, since we only observe allele frequencies and not haplotypes, we do not see recurrent mutations of the same type as separate segregating sites. We would consequently under-count recurrent mutations on autosomes, and may observe an apparent enrichment of these types on the X-chromosome. For this reason, differences in the relative diversity at CpG sites on the X chromosome and autosomes must be cautiously interpreted. This concern applies not just to CpG transitions (C>T mutations at CpG sites), which have the highest mutation rate, but potentially also to C>A and C>G mutations at CpG sites, which also have a higher mutation rate than average (7, 8), and for which we observed a substantial decrease in X-enrichment when we counted multiple alleles at a site (Supplementary Figure 3). To be conservative, we excluded CpG sites in comparisons between the X chromosome and autosomes. Including them does not change any of our qualitative results, however. The explanation is likely that the difference in effective population size for the X chromosome and autosomes is small, and that the effect of recurrent mutations on the X-autosome comparison is even smaller. This minor effect is mitigated further by including visible multi-allelic sites.

**Testing for significant differences in the mutation spectrum between genomic compartments.**

We tested if mutation type $i$ is distributed the same way in two compartments (a and b) given what would be expected based on the overall distribution of mutations in the two compartments. Effectively, we considered the relationship between the following four ratios for each of 96 mutation types (excluding the 16 CpG types where needed):

| | |
|---|---|
| $\pi_{i(a)}$ | $\pi_{i(b)}$ |
| $\pi_{(a)}$ | $\pi_{(b)}$ |

Or

| | |
|---|---|
| $(n_{i(a)}/d_{i(a)})$ | $(n_{i(b)}/d_{i(b)})$ |
| $(n_a/d_a)$ | $(n_b/d_b)$ |

We designated the larger compartment (i.e., with a greater number of mutational opportunities) as the reference compartment (for example, in X-autosome comparisons, the autosomes were used as reference). We assumed that the number of mutations of a particular type in compartment a (the "test" compartment) is binomially distributed with a mean value proportional to the observed diversity for that type in compartment b (the reference compartment), adjusted for overall differences in diversity between the two compartments:

$$n_{i(a)} \sim \text{binom}(d_{i(a)}, f_i)$$

$$E(n_{i(a)}) = d_{i(a)} * f_i$$

$$\text{where } f_i = \pi_{i(b)} * \frac{\pi_{(a)}}{\pi_{(b)}} = (n_{i(b)}/d_{i(b)}) * \frac{(n_{(a)}/d_{(a)})}{(n_{(b)}/d_{(b)})}$$

The factor $f_i$ is the expected diversity level for a given type in the test compartment. For each type, we tested if the observed number of mutations in the test compartment differs from the number expected by chance, using the "binom.test" function in R to obtain p-values. Mutational types were considered significantly different in their frequencies in the two compartments if the two-tailed p-value from the binomial test was below the Bonferroni-corrected 5% significance threshold (=0.05/96). The relative enrichment for each mutation type is given by:

$$\frac{n_{i(a)}}{E(n_{i(a)})} = R_{i(ab)}$$

We also obtained 95% confidence intervals for the relative enrichment of each type from the binomial test.

We implicitly assumed that mutations of one type do not impact mutational opportunities of other types. The reason is that because the total number of mutations is much smaller than the number of mutational opportunities, an increase in the number of mutations of one type does not appreciably decrease the mutational opportunities available for other types. We note, however, that the tests for different mutation types are still not fully independent, because the expected diversity for each mutation type in the test compartment depends on the overall relative diversity in the two compartments. If they constitute a large proportion of the total number of mutations, mutation types that are highly significantly enriched in one compartment could influence the null distribution for other mutation types and thus lead to the depletion of these other types in that compartment. In general, we focused on describing the top signals we observed, which are unlikely to be strongly affected by this phenomenon.

Nevertheless, to assess the impact of this issue, we implemented a procedure similar to the "forward variable selection procedure" used by Harris and Pritchard (2017). We ranked mutation types by their p-values in an initial set of 96 tests. We then sequentially removed the most significant mutation type and reassessed the other types for significance at each step; mutation types that reached significance through interactions with other types should drop out. We note that re-generating the ratio of expected diversity in two genomic compartments at each step based on the mutation types remaining in the sample can result in even more significant differences between compartments. Because mutations at CpG sites have the largest sample sizes by far, the largest impact of forward variable selection is observed when CpG transitions are highly enriched in a particular compartment (i.e., in non-CpG islands relative to CpG islands); for this analysis we only highlight the top signals (Supplementary Fig. 11a). Our other analyses remain qualitatively unchanged by forward variable selection, and also by excluding CpG sites. The effect of these procedures on our X-autosome comparison is shown in Supplementary Figure 4.

We note that in testing for significant differences in the mutation spectrum between genomic compartments using a binomial test, as described above, we implicitly ignore sampling error in the estimate of diversity of the designated reference compartment; we verified that our results are insensitive to this assumption by using alternative approaches to calculate significance that do not make this assumption, but have other limitations. These are described below:

First, we bootstrapped the expected distribution of a particular mutation type in the two compartments using hypergeometric sampling. For a given mutation type $i$ with sample size $n_i(= n_{i(a)} + n_{i(b)})$, we generated a random variable $k_i$ for the number of draws in compartment a when $n_i$ mutations were drawn without replacement from the pool of all mutations in both compartments, $n = (n_{(a)} + n_{(b)})$ with $n_{(a)}$ "successes" in compartment a, i.e., $k_i = Hyper\big(n, n_{(a)}, n_i\big)$. The expected distribution of the ratio of mutations of type $i$ in the two compartments, $k_i/(n_i$-$k_i)$, was obtained using 10,000 such trials. We calculated a p-value using the rank of the observed relative diversity (adjusted by the ratio of total mutational opportunities of all types in the two compartments) in the expected distribution. In this approach, we allowed for uncertainty in the number of mutations of a particular type in both compartments, but held constant the total diversity in each compartment.

Second, we adjusted the number of mutations in one of the compartments by the ratio of overall diversity in the two compartments, and then applied a chi-squared test to the 2x2 contingency table (shown below) of the mutations ("successes") and the remaining mutational opportunities ("failures") by compartment. Third, we fitted the same 2x2 contingency table using a binomial glm (with the compartment as a covariate).

| | |
|---|---|
| $n_{i(a)} * 1/p$ | $n_{i(b)}$ |
| $d_{i(a)} - (n_{i(a)} * 1/p)$ | $d_{i(b)} - n_{i(b)}$ |

$$where\ p = (n_a/d_a)\,/\,(n_b/d_b)$$

We compared these methods for three analyses: X vs Autosome, PAR vs Autosome, and X-A matched for replication timing (Supplementary Methods Tables 1-3). In all cases, the same significant mutation types (and the same effect sizes) stand out; in other words, the results are the same, regardless of the approach. The reason is likely that the reference compartment is always sufficiently large for sampling error to be small.

**Comparing the X chromosome and autosomes: additional considerations.**

In comparing compartments on the X chromosome and autosomes, we excluded the pseudo-autosomal region unless otherwise specified, since sex-specific properties differ between the PAR and the rest of the X chromosome.

We considered additional population-level properties that may affect the mutation spectrum differently for the X and autosomes. In accordance with our prior expectation that biased gene conversion should have a negligible effect on variants at very low frequencies, we note no clear patterns of X-autosome differences for mutation types that are subject to biased gene conversion. Similarly, because so many variants in the sample are rare and thus young, we expect very little effect of either direct or linked selection on the mutation spectrum a priori. Thus, we interpret these mutation patterns as reflecting real and largely neutral differences between X and autosomes.

We also checked for differences in genotyping error rates between the X chromosome and autosomes. Because the variant quality (QUAL) variable in the dataset is jointly calculated based on males and females in the sample, the reported quality of variants on the X chromosome is expected to be slightly lower. Nevertheless, the distribution of variant quality is almost identical for the X chromosome and autosome (Supplementary Figure 6), and any small differences are likely further lessened because we used only the female subset of the data. To rule out a potential interaction of error rate by mutation type and compartment, we compared the genotype qualities and read depths for C>G and C>A mutation types in compartments across which we found the distribution of these types to differ. The average genotype quality and read depth is high for all mutation types, and while there may potentially be small differences in error rates between different types of mutation, these do not seem to differ across compartments in ways that would affect our inference. For instance, average genotype quality and read depth for C>G mutations is similarly distributed to all other mutation types combined both in the PAR and Autosomes, while a strong enrichment of these types is seen only in the PAR. We similarly did not observe a notable interaction of C>A quality metrics with early and late replicating compartments; these cases are illustrated in Supplementary Figure 7.

We further replicated our analysis using two datasets (Uk10k and SGDP) (The UK10K Consortium, 2015; Mallick *et al.*, 2016) sequenced independently, with varying levels of coverage (Supplementary Fig. 5).

In order to validate signals observed in the gnomAD data, we conducted a similar analysis using publicly available data from the Simons Genome Diversity Project (SGDP) (10). We considered only individuals with ID beginning with "S" (as this subset was sequenced PCR-free and processed using a consistent approach) and used filter level 1 (recommended as optimal for SNP discovery in Mallick et al. 2016) to obtain a variant set of high quality. The SGDP variants were polarized with respect to the major allele in the full SGDP sample of "S" individuals. We limited our analysis to female individuals (100/256 individuals in the dataset). Individual-level alternate allele counts at each variant position, reported as 0, 1 (heterozygous), or 2 (homozygous), were summed over the 100 individuals to obtain allele counts in the sample; no multi-allelic sites were seen in this sample. Sites with >50% missing data were excluded. The major allele in the SGDP dataset is 99% correlated with the gnomAD major allele when only variants at matched positions are considered. To estimate the mutational opportunities for each type of site in this dataset, we combined accessible regions from 11 individuals (five with predominantly European and six with African ancestry).

We also replicated our analysis using variants in the ALSPAC and TwinsUK subsets of the UK10K dataset (The UK10K Consortium, 2015). We limited our analysis to the 2,793 females. We polarized to the major allele in the UK10K sample, and applied quality thresholds similar to those in the gnomAD analysis. We excluded multi-allelic sites from this sample. Since accessible regions were unavailable for this dataset, we used the accessible regions obtained for the SGDP dataset, obtained as described above.

**Obtaining data for the distribution of genomic features.**

To investigate the association of biochemical features with mutational patterns in the germline, we would ideally consider the distribution of these features in germline tissue. In cases where we were limited to data from somatic cell lines, where possible we focused on genomic features that are known to have stable or broadly similar distributions across tissues-types, as these are more likely to be comparable between the soma and germline.

We downloaded replication timing data from two sources: data for LCL lines was obtained from Koren et al., 2012, and data for three human embryonic stem cell lines (H1, H7, and H9), produced as part of the ENCODE project, was downloaded from the UCSC browser. Replication timing data are reported as a standardized score with negative scores representing later replication. To check for systematic differences in the observed distributions of replication timing between the two studies, we also downloaded data for the cell line most similar to LCL available from ENCODE (GM12878); for these cells, the distribution of replication timing was almost identical to the LCL data from Koren et al., 2012, suggesting that differences in the distributions of replication timing between the LCL and embryonic stem cell lines are not due to methodological differences, and likely reflect real biological differences between cell lines. For autosomes, average replication timing is largely consistent across different cell lines, both at the chromosomal level and at the 1Mb scale (Supplementary Figure 8) (11, 12).

Since methylation is expected to be highly variable across tissues and may well differ substantially between the germline and soma, we used CpG islands as a binary proxy for methylation: CpG islands are hypomethylated relative to the rest of the genome, across tissue-types (13, 14).

We obtained the X-inactivation status ("inactive", "escape", "variable", "unknown") of genic regions on the X chromosome from Tukiainen et al. 2017; this consensus status for 683 genes on the X chromosome is based on combined information from multiple sources and experimental approaches, across tissue-types. To be consistent with the Tukiainen et al. 2017 study, we used Gencode v19 coordinates and annotations for genic regions on the X chromosome and autosomes. The small number of regions of overlap between genes that were classified as both "escape" and "inactive" were excluded from the analysis.

DMC1 ChIP-seq signal intensity on the X chromosome and autosomes, measured in spermatocytes, was obtained from Pratto et al., 2014. DMC1 hotspots were defined as 1 kb regions around the midpoint of hotspots identified by Pratto et al., 2014. We used hotspots and signal intensity values for the "AA2" individual; using average intensities and the union of hotspots from all four individuals with PRDM9 alleles A and B does not alter our qualitative results. DMC1 hotspots were grouped as weak (DMC1 signal intensity 1-50), intermediate (signal intensity 50-150), and strong (signal intensity >150). Because the X chromosome has a disproportionate number of very strong DMC1 hotspots, we chose these criteria to obtain similar average hotspot intensities on the X chromosome and autosomes in the first three bins, with the outliers in the fourth; varying these thresholds does not alter our qualitative conclusions. We obtained the list of autosomal regions enriched for clustered C>G mutations from Jónsson et al., 2017. Finally, we used the female standardized recombination map (16) to define female hotspots on autosomes (following Kong et al. 2010, windows with recombination rates greater than 10-fold the genome average were considered hotspots; increasing this threshold does not alter our qualitative conclusions).

**Testing the effect of replication timing and other genomic features on the autosomal mutation spectrum.**

For autosomes, average replication timing is largely consistent across cell lines, both at the chromosomal level and at the 1Mb scale (Supplementary Figure 8) (11, 12). We compared the mutation spectrum in autosomal regions that are early or late replicating in LCL (Fig. 2a) and H9-hESC (Supplementary Fig. 9a) cell lines. Regions were defined as early replicating if the replication timing score was greater than or equal to 0.5, and late replicating if it was less than or equal to -0.5 (the results remain qualitatively unchanged if these thresholds are varied).

We aggregated replication timing data per 1 Mb window and per chromosome (using the bedtools map function). While the choice of scale is somewhat arbitrary, averaging replication timing on the 1 Mb scale is relatively lossless (Supplementary Figure 8), and this scale has been used in other studies of replication timing (Smith, Arndt, and Eyre-Walker 2018; Hiratani et al. 2010). In each 1 Mb window, we obtained the enrichment of each of six broad mutational classes (C>A, C>G, C>T, T>A, T>C, T>G) relative to all mutation types, and relative to all windows taken together. We excluded a small number of windows in which the total number of mutations was outside the range of two standard deviations from the mean. For the chromosome level analysis, for each autosome, we obtained the enrichment of each of the six broad mutational classes relative to all mutation types, and relative to all other autosomes taken together. Averaging replication timing on entire chromosomes is useful because it allows us to place the effect of replication timing on the X chromosome in context.

To assess the impact of other genomic features, we compared the autosomal mutation spectrum in regions that lie within and outside CpG islands, and regions that lie in regions of clustered de novo mutations thought to be due to double-strand break damage (Supplementary Figure 11b). We note that many genomic features are correlated. As one example, hypomethylated CpG islands tend to colocalize with early replicating gene regulatory regions; we consider the effect of this interaction on the mutation spectrum (Supplementary Figure 10).

**Controlling for genomic features on the X-chromosome.**

For the X chromosome, the average difference in replication timing between cell lines is thought to reflect X-inactivation status, which differs by cellular genotype and the level of differentiation, and can be heterogenous in a cell population (11, 12, 18–20). For instance, the LCL line is a female (XX) cell line with one stably inactivated X chromosome, consistent with the significantly later replication of the X on average (Supplementary Figure 8). The H1 embryonic cell line is male (XY), whereas the H7 line is XX but with two active X chromosomes, and in both these cases the X replicates on average at the same time as autosomes (Supplementary Figure 8). We did not observe late replication on the X chromosome for the female H9 cell line (Supplementary Figure 8), which is thought to have one inactive X chromosome. The reason may be that this particular line is derived from 5-day old female embryos: since the pre-implantation embryo undergoes global demethylation and is hypomethylated around day 5 post-fertilization (21, 22), we do not necessarily expect to see an effect of X-inactivation in this cell line. Thus, we did not have data for human embryonic stem cells where one X chromosome is stably inactive; however, the differences in X-inactivation for human embryonic cells in various states of differentiation is consistent with X-inactivation status changing over time in the germline, and on average lying somewhere in between that observed in the LCL cell lines and the human embryonic stem cell lines. More generally, these patterns support the notion that X-inactivation and replication timing on the X chromosome are highly correlated.

We controlled for X-inactivation on the transcribed X chromosome using the X-inactivation status. Complementary to this approach, and to ensure that the observed impact of X-inactivation was not an artifact of a potential mis-classification of genes in the active and inactive categories, we controlled for replication timing on the X chromosome. To match the average replication timing on the X and autosomes, we considered regions on the X chromosome and autosomes that have replication timing scores of greater than or equal to -0.5 and less than or equal to 0.5 in the LCL cell line (Supplementary Fig. 13). The total length of callable regions (in gnomAD) with this property is about 700 Mb on autosomes and 50 Mb on the X. On average, variants in these regions on both the X and autosomes have a similar (approximately zero) mean score for replication timing. In matching the mean, we implicitly assumed that the effects of replication timing on the mutation spectrum were roughly linear (there are some non-linear effects on the X chromosome in regions of extremely late replication timing). Changing the threshold does not alter our qualitative results as long as the mean replication timing on the X chromosome and autosomes is close.

We also considered shared mutational patterns in early and late replicating regions of the X relative to autosomes (Supplementary Fig. 14). Since the mean replication timing on the X is -0.75, we considered late replicating regions to be those with a replication timing score < -1.25, and early replicating regions, > -0.25; changing these thresholds changes the power of our comparison but not the qualitative results.

We ignored effects of differences in CpG methylation between the X and autosome since we excluded CpG sites in the X-autosome comparison. We assumed that any additional small effects of methylation at other sites were controlled for indirectly by controlling for replication timing and/or X-inactivation.

For the pseudoautosomal compartment, we did not control for genomic features, since it does not undergo inactivation and replicates early and thus, the mutation spectrum in this compartment should not be affected by these features.

### Supplementary Figures

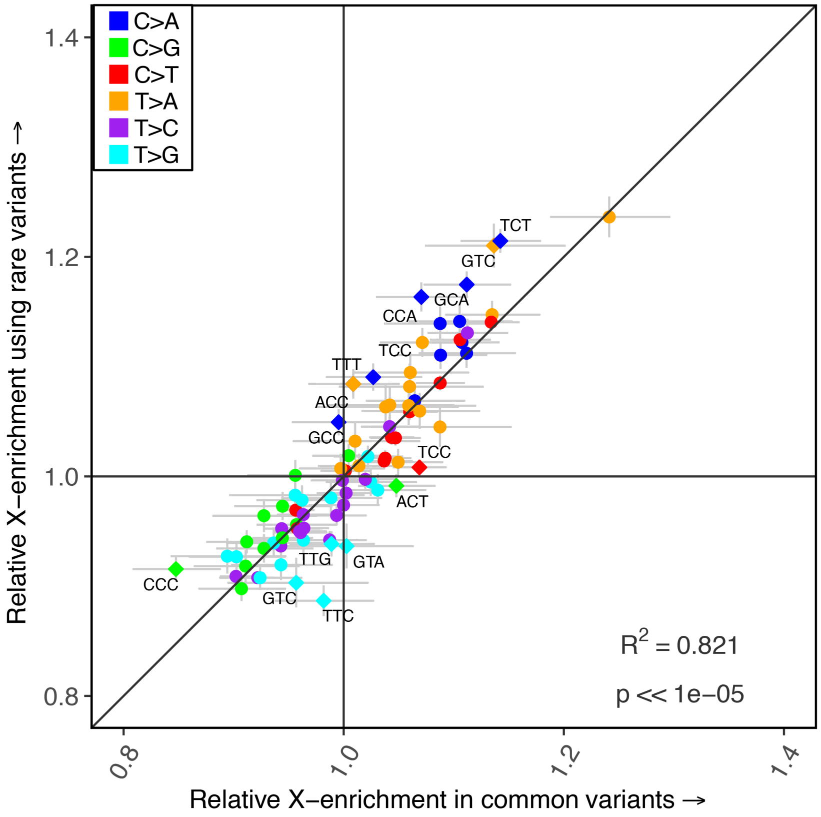


**Fig. S1.** The enrichment of mutation types on the X chromosome, relative to autosomes, for rare variants (allele counts 1-5) versus common variants (variants at frequency 1% or greater) in the sample. This analysis excludes CpG sites and the pseudoautosomal region. Diamonds represent mutation types that differ by 5% in their mean fold enrichment between the two comparisons. The black diagonal represents the x=y line. For each estimate, the 95% confidence interval from a binomial test for X-enrichment among common and rare variants, respectively, is represented by the grey horizontal and vertical bars.

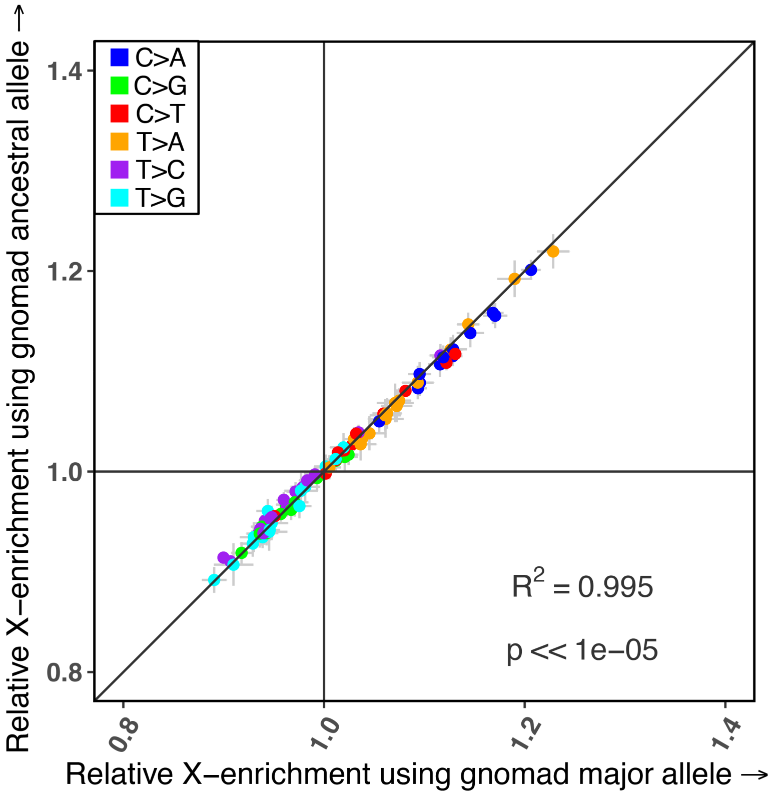


**Fig. S2.** The enrichment of mutation types on the X chromosome, relative to autosomes, using the major allele in the sample versus the ancestral allele from the human chimp ancestral sequence. This analysis excludes CpG sites and multi-allelic sites. The black diagonal represents the x=y line. For each estimate, the 95% confidence interval from a binomial test for enrichment on the X chromosome relative to autosomes, using the major allele or the human chimp ancestral allele, respectively, is represented by the grey horizontal and vertical bars.

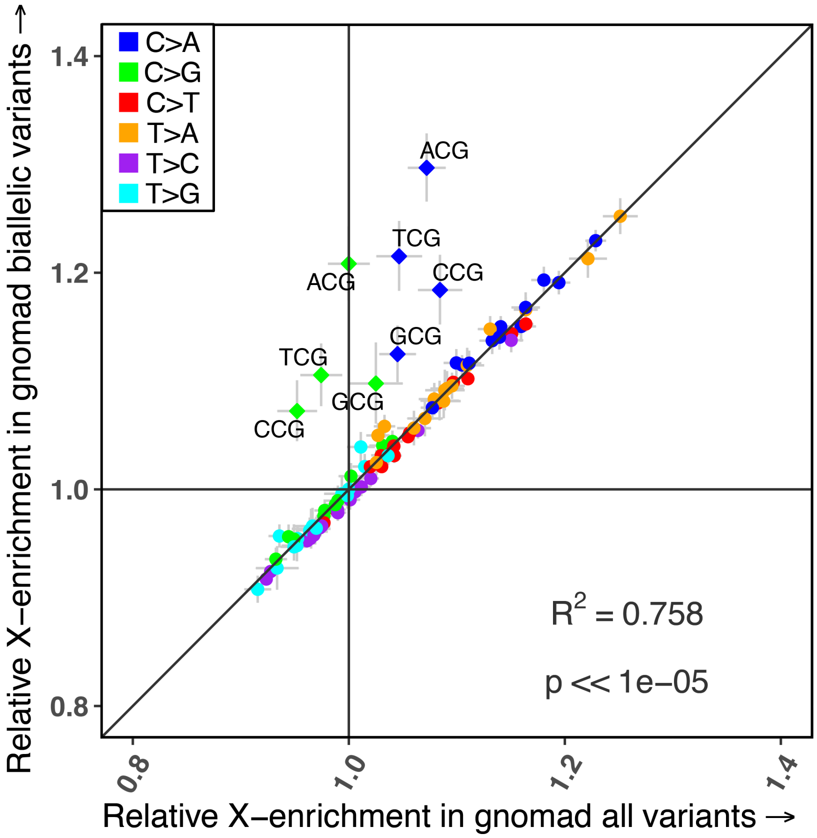


**Fig. S3.** The enrichment of mutation types on the X chromosome, relative to autosomes, in all variants including multi-allelic sites, or bi-allelic sites only. Diamonds represent mutation types that differ by 5% in their mean fold enrichment between the two comparisons. The black diagonal represents the x=y line. For each estimate, the 95% confidence interval from a binomial test for enrichment on the X chromosome relative to autosomes, using all sites or only bi-allelic sites, respectively, is represented by the grey horizontal and vertical bars.

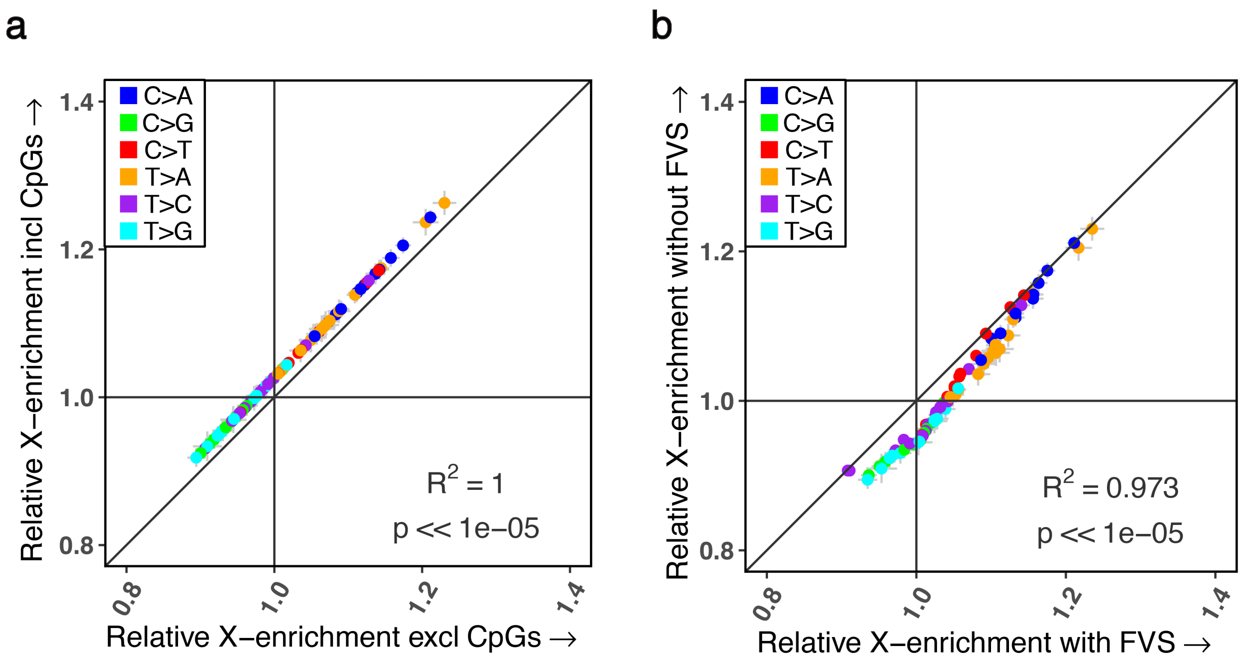


**Fig. S4.** The effect of forward variable selection and excluding CpG sites on the X-autosome comparison. The black diagonal represents the x=y line. For each estimate, the 95% binomial confidence intervals are represented by the grey horizontal and vertical bars. (a) The enrichment of mutation types on the X chromosome, relative to autosomes, when all 96 types are considered, versus when CpG sites are excluded. (b) The enrichment of mutation types on the X chromosome, relative to autosomes, with and without the forward variable selection procedure (see Supplementary Methods).

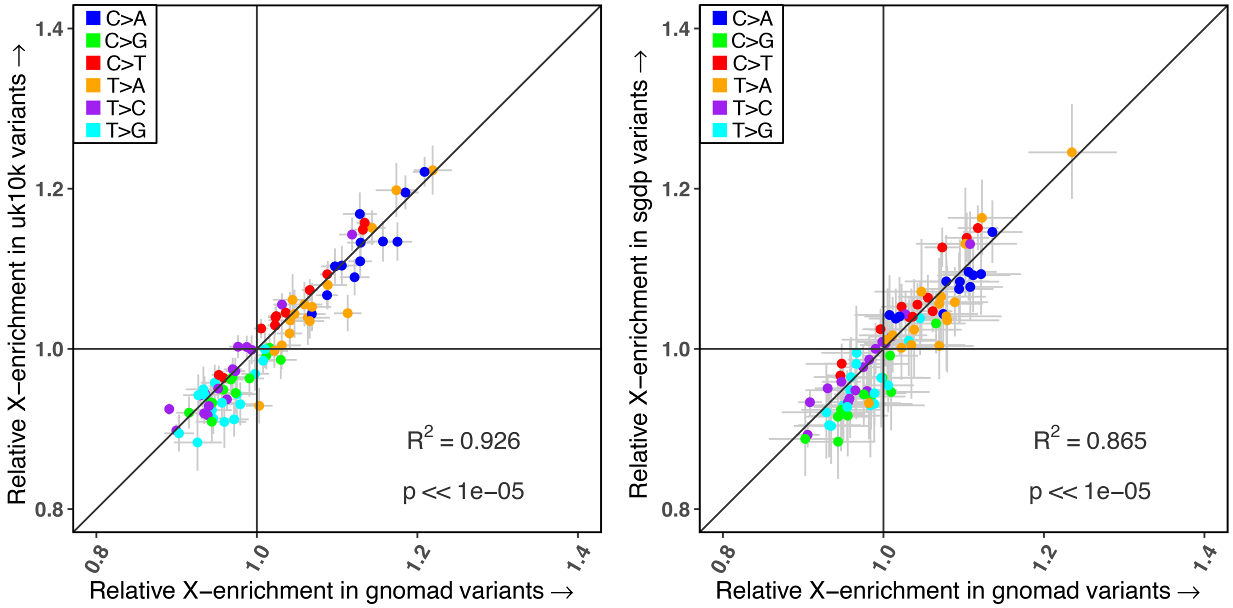


**Fig. S5.** Comparison of the X-Autosome mutation spectrum in gnomAD with the UK10K and SGDP datasets. The gnomAD dataset was down-sampled to match the comparison dataset in each case. These analyses exclude CpG sites and multi-allelic sites. The black diagonal represents the x=y line. For each estimate, the 95% confidence interval from a binomial test for X-enrichment relative to autosomes in gnomAD and the UK10K or SGDP datasets, respectively, is represented by the grey horizontal and vertical bars. (a) The enrichment of mutation types on the X chromosome, relative to autosomes, in gnomAD versus UK10K. (b) The enrichment of mutation types on the X chromosome, relative to autosomes, in gnomAD versus SGDP.

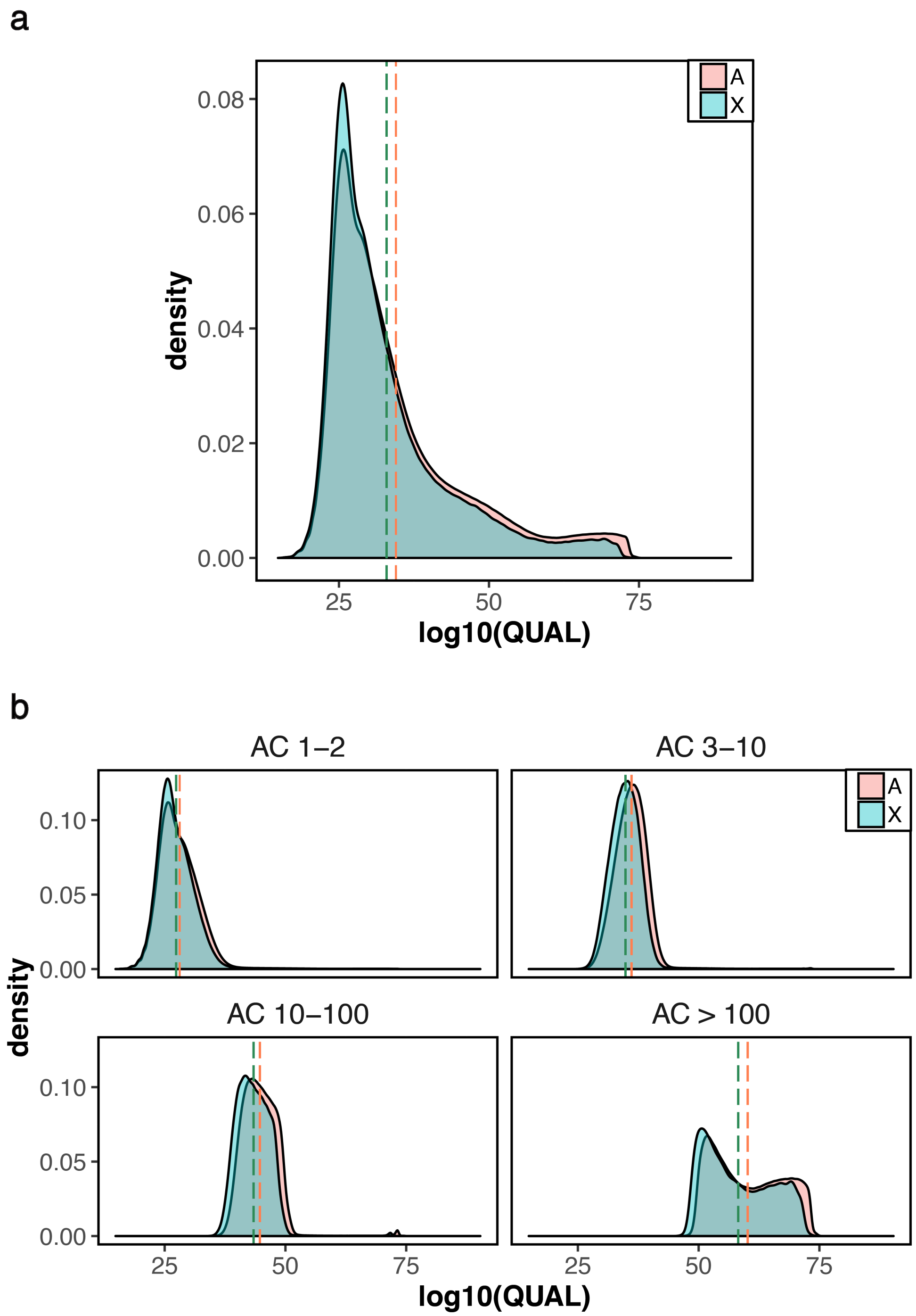


**Fig. S6.** (a) Distribution of Variant Quality on the X chromosome and Autosomes (b) Distribution of Variant Quality on the X chromosome and Autosomes by frequency bin. Note that this reported measure of variant quality is based on the full sample with males and females and might be expected to be slightly lower on the X chromosome (see Supplementary Methods).

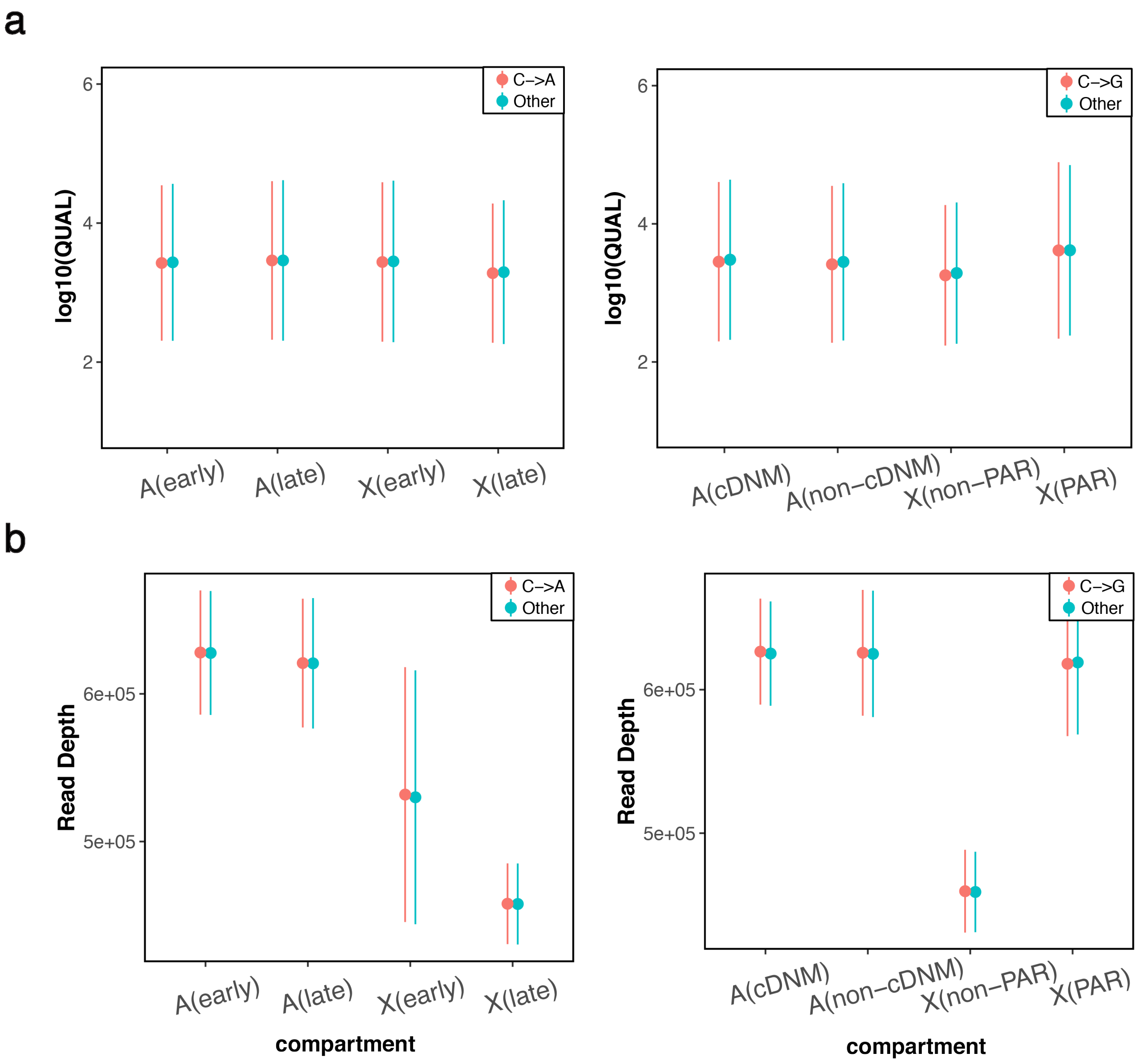


**Fig. S7.** Average genotype quality and read depth by mutation type and compartment. Note that these reported measures of variant quality are based on the full sample with males and females and might be expected to be slightly lower on the X chromosome. (a) Genotype quality for C>A mutations types in X-Autosome compartments with differences in replication timing, and for C>G versus all other mutation types in the PAR and other relevant X-Autosome compartments. (b) Read Depth for C>A mutations types in X-Autosome compartments with differences in replication timing, and for C>G versus all other mutation types in the PAR and other relevant X-Autosome compartments.

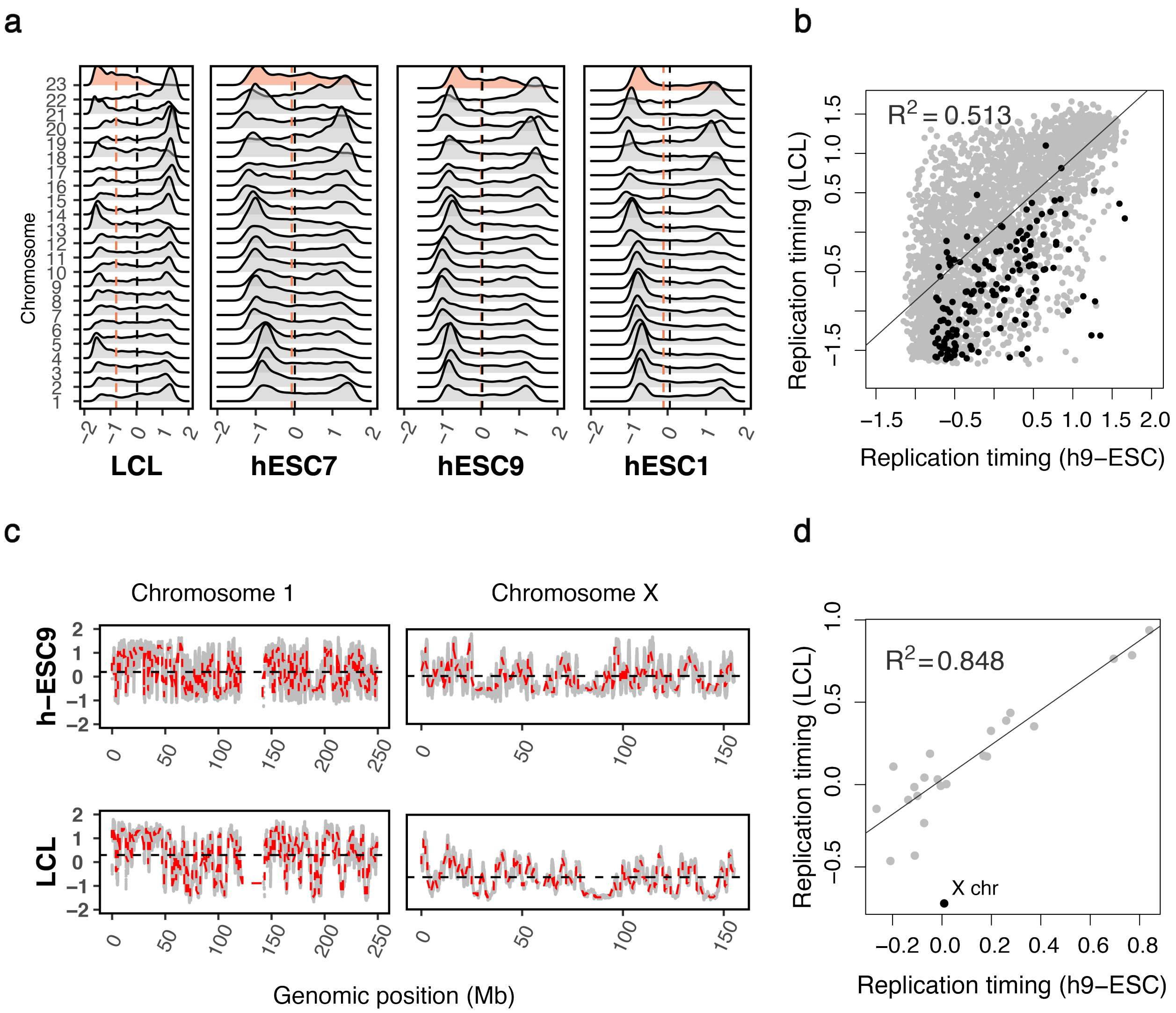


**Fig. S8.** Variation in replication timing scores by cell type. Positive values indicate early replication. (a) The distribution of replication timing scores for LCL and human embryonic stem cell lines (hESC1, hESC7, hESC9). The X chromosome is shaded in red. The average scores on autosomes and on the X are denoted by the red and black vertical lines, respectively. (b) Mean replication timing for 1 Mb windows for the LCL and hESC9 cell types. Autosomal windows are shown in grey; windows on the X chromosome have been overlaid in black. (c) Fine scale replication timing for chromosomes 1 and X. Grey points reflect raw replication timing data in bins of approximately 100 bp. The dashed red lines reflect averages over 1 Mb windows. The horizontal black line indicates the chromosome-level mean replication timing. (d) Mean replication timing at the chromosomal level for the LCL and hESC9 cell types.

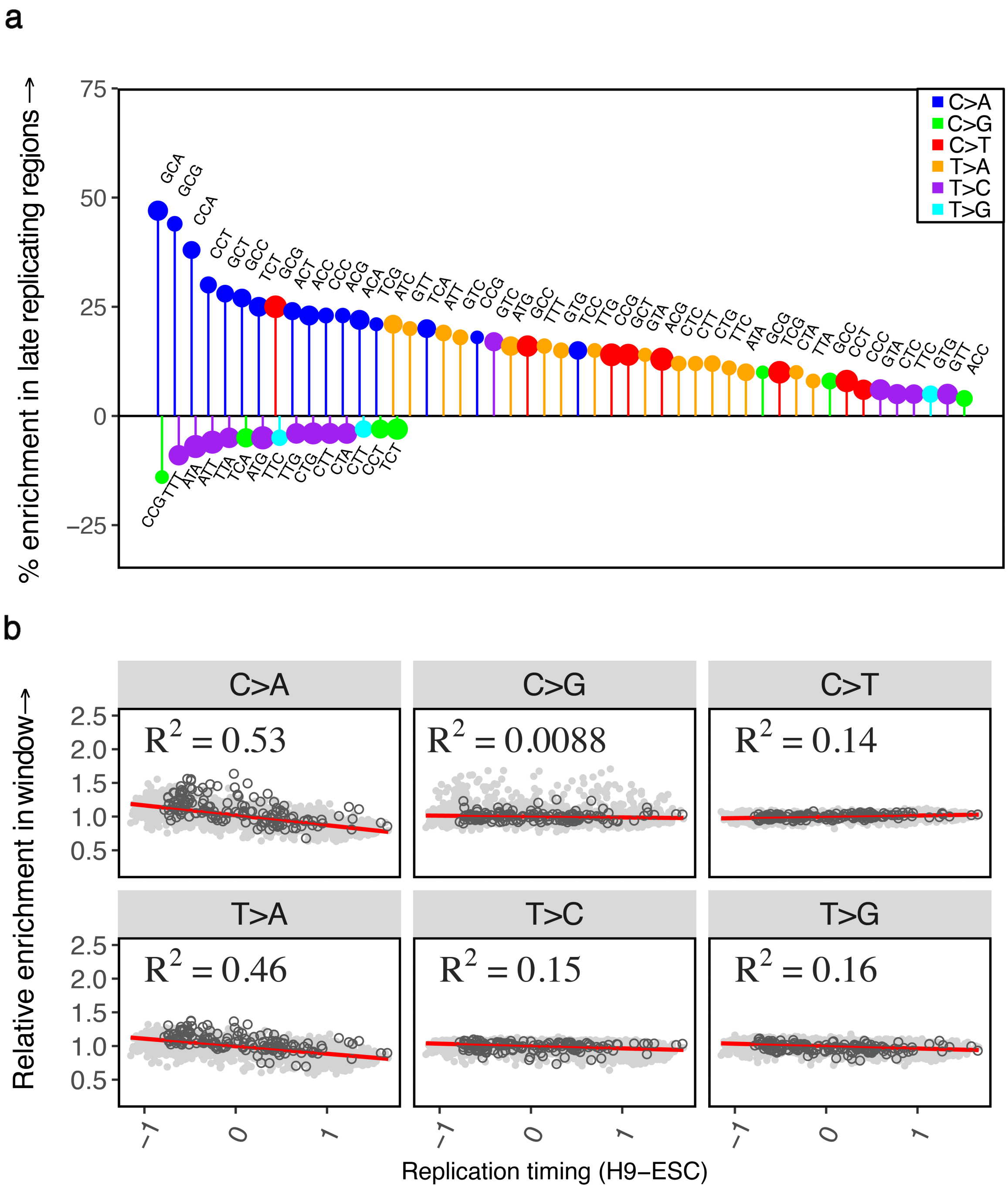


**Fig. S9.** The effect of replication timing on the mutation spectrum using the hESC9 cell type. (a) Comparison of the spectrum of 96 mutation types in late replicating autosomal regions relative to early replicating autosomal regions. Only significant differences are shown. Positive and negative effects are ranked separately in order of effect size from left to right. Late replicating regions are defined as having a replication timing score <= -0.5 and early >= 0.5. (b) The relative enrichment of six mutational classes in 1Mb windows relative to all autosomal windows combined, ordered by the mean replication timing in 1Mb windows. Positive x-values indicate early replication. Windows on autosomes are shown in solid grey circles; windows on the X chromosome have been overlaid in black open circles.

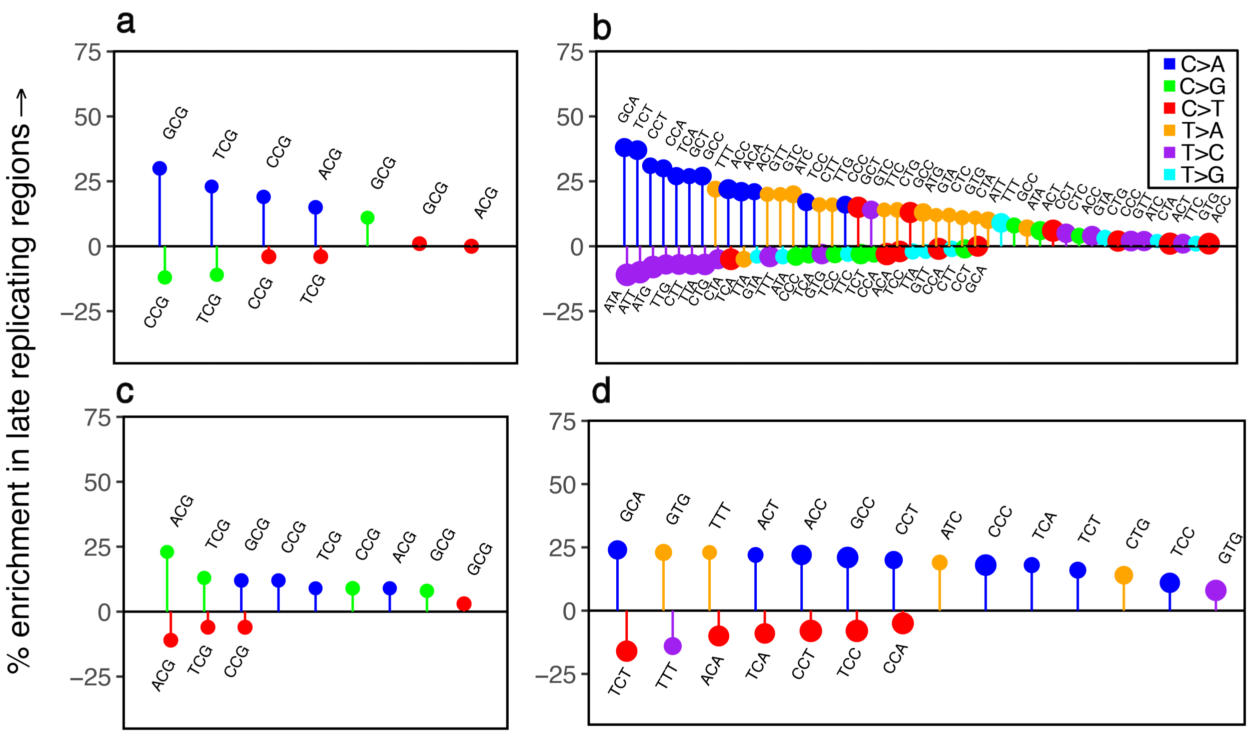


**Fig. S10.** The effects of methylation and replication timing on the mutation spectrum. Only significant differences are shown. Positive and negative effects are ranked separately in order of effect size from left to right. CpG islands are used as a proxy for regions of hypomethylation. Late replicating regions are defined as having a replication timing score <= -0.5 and early >= 0.5. (a) Comparison of the mutation spectrum at CpG sites in late versus early replicating regions outside CpG islands. (b) Comparison of the mutation spectrum at non-CpG sites in late versus early replicating regions outside CpG islands. (c) Comparison of the mutation spectrum at CpG sites in late versus early replicating regions inside CpG islands. (d) Comparison of the mutation spectrum at non-CpG sites in late versus early replicating regions inside CpG islands.

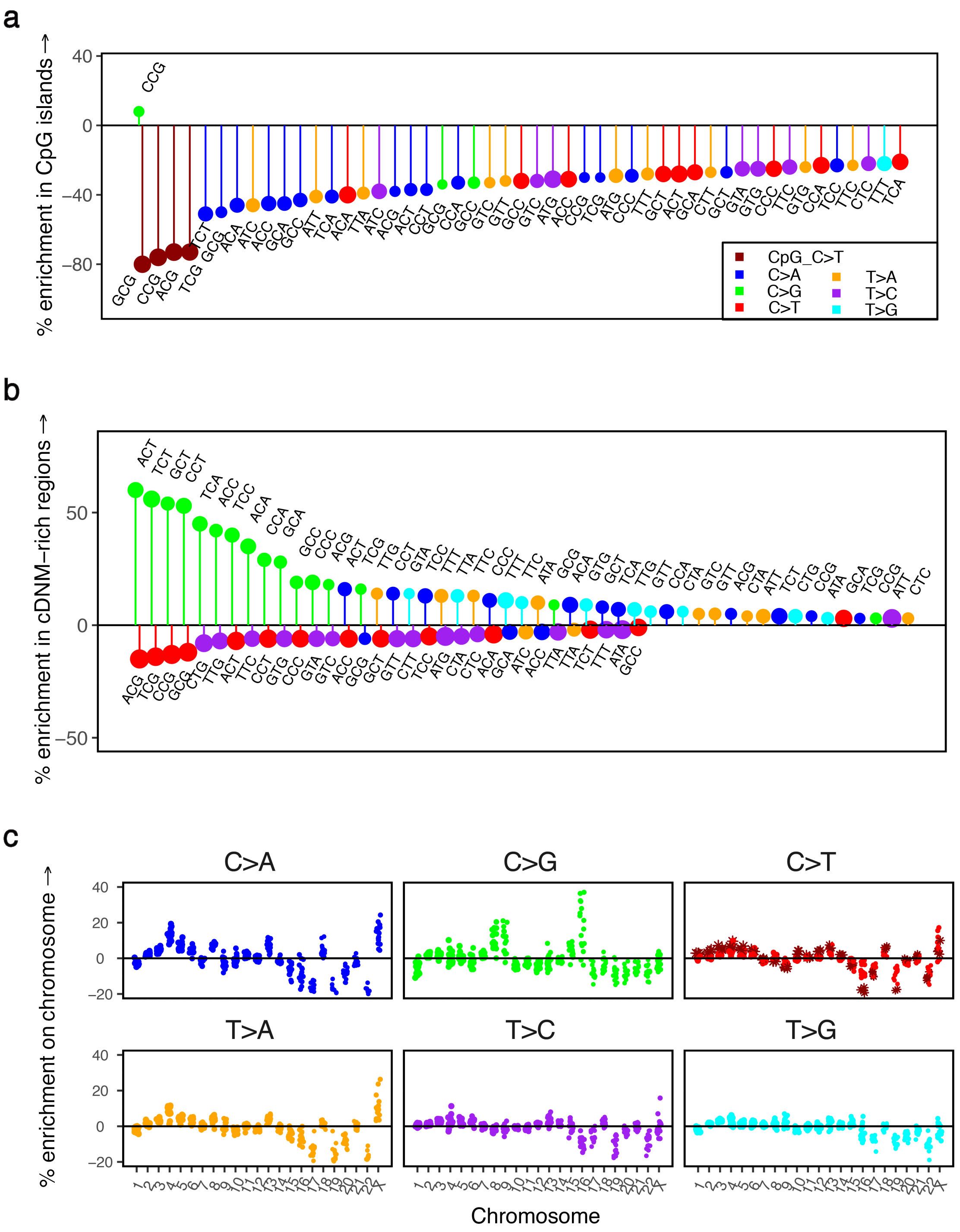


**Fig. S11.** The effect of biochemical features on the mutation spectrum. (a) Comparison of the spectrum of 96 mutation types in autosomal regions in CpG islands relative to autosomal regions outside CpG islands. Positive and negative effects are ranked separately in order of effect size from left to right; only the top 50 significant positive and negative effects are shown for legibility. The size of the circle reflects the number of mutations of that type. CpG transitions are labeled in dark red. (b) Comparison of the spectrum of 96 mutation types in autosomal regions identified as rich in clustered de novo mutations (cDNMs) by Jónsson et al., 2017, relative to other autosomal regions. Only significant differences are shown. Positive and negative effects are ranked separately in order of effect size from left to right. The size of the circle reflects the number of mutations of that type. (c) Variation in the mutation spectrum for individual chromosomes. The enrichment level is shown for individual autosomes relative to all other autosomes combined and, on the X, relative to all autosomes combined; each point reflects one of 16 trinucleotide contexts for a particular mutational class. CpG transitions are labeled in dark red.

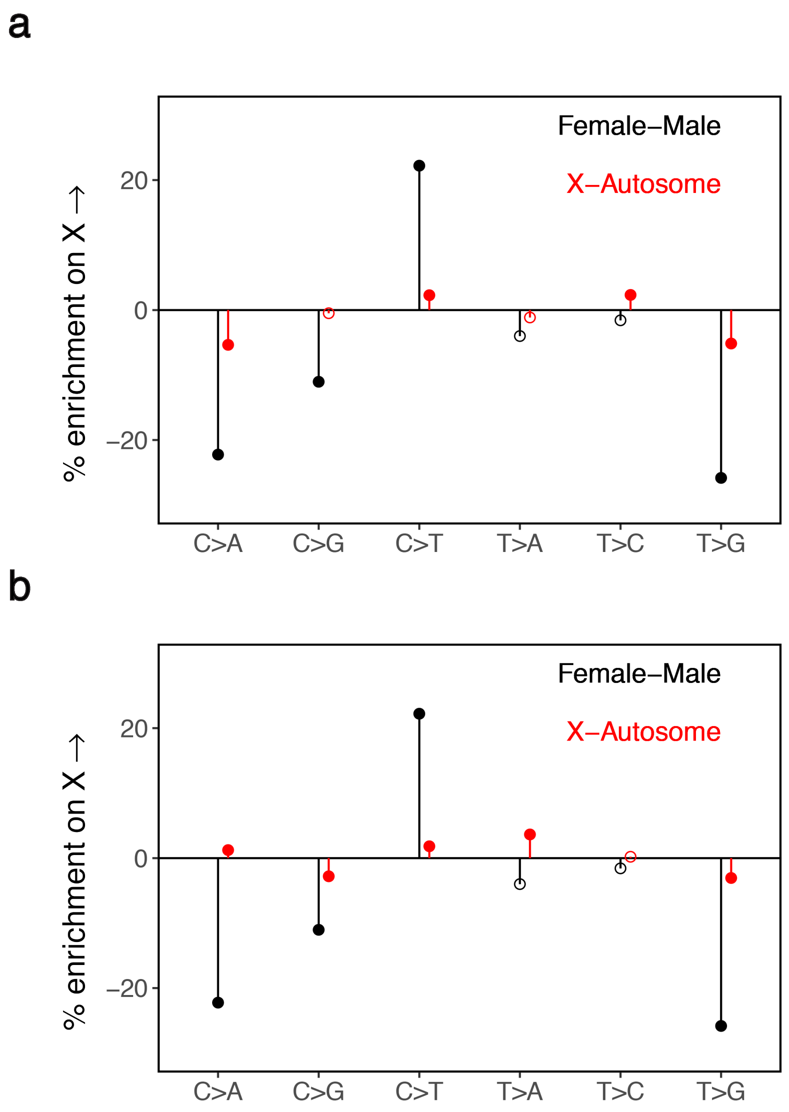


**Fig. S12.** The mutation spectrum in the active and inactive genic regions of the X chromosome relative to genic regions in autosomes. These X-Autosome comparisons exclude all CpG sites and the pseudoautosomal region (PAR). (a) The spectrum of six mutational classes in the active genic regions of the X chromosome relative to autosomes (in red), compared to known male female differences from Jónsson et al., 2017 (in black). Solid points are statistically significant differences at the 5% level, accounting for multiple tests. (b) The X-Autosome mutation spectrum in inactive genic regions of the X relative to autosomes (in red) compared to known male female differences from Jónsson et al., 2017 (in black). Solid points are statistically significant differences at the 5% level, accounting for multiple tests.

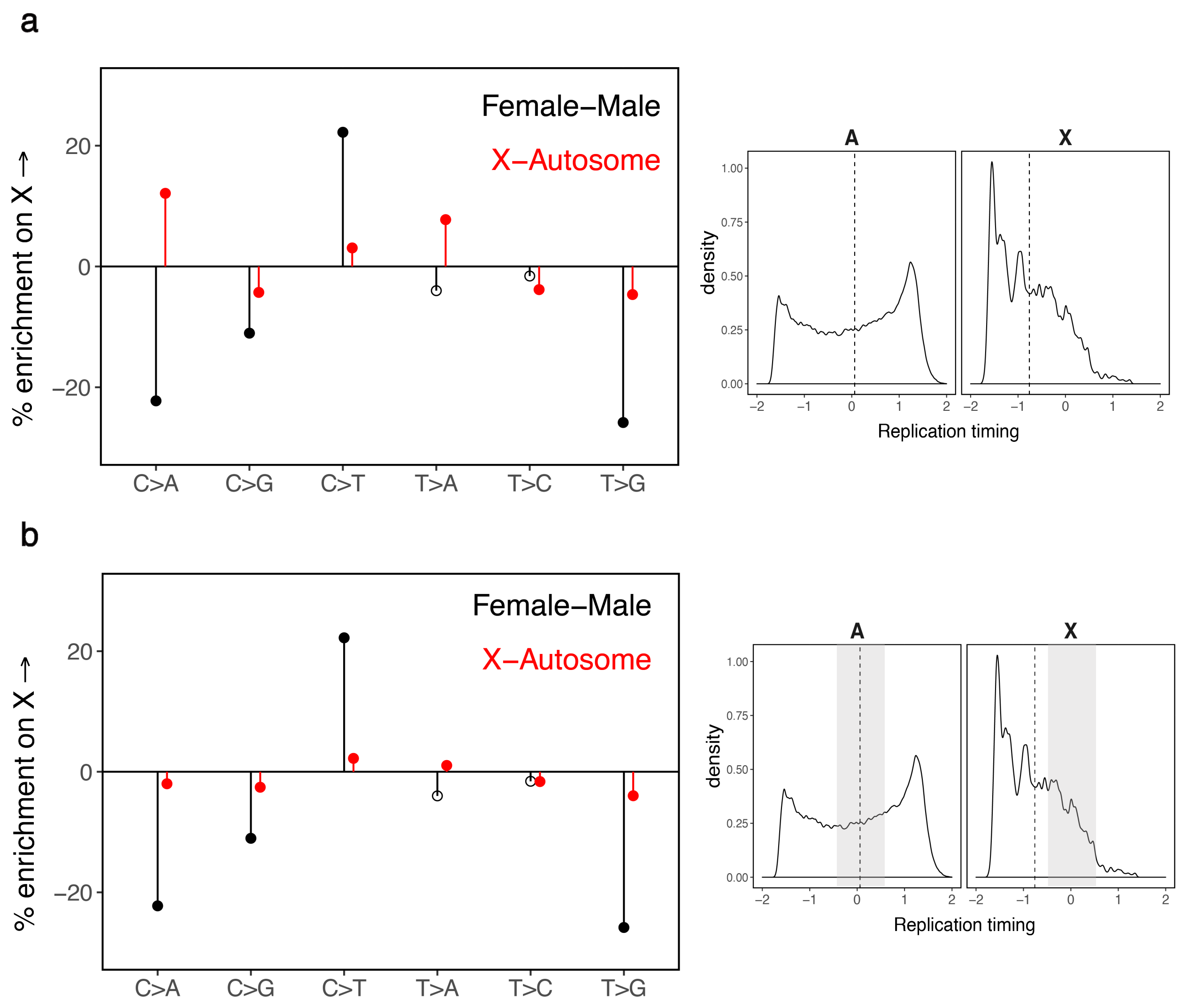


**Fig. S13.** The mutation spectrum on the X chromosomes and autosomes with and without matching for average replication timing. These X-Autosome comparisons exclude all CpG sites and the pseudoautosomal region (PAR). The distribution of replication timing for the X and autosomes is shown in the right panels, with the mean replication timing on the X and autosomes represented by dashed vertical black lines. (a) The X-Autosome mutation spectrum (in red), unadjusted for replication timing differences, compared to known male female differences from Jónsson et al., 2017 (in black). Solid points are statistically significant differences at the 5% level, accounting for multiple tests. (b) The X-Autosome mutation spectrum of the X and autosome matched for average replication timing, in red, compared to known male female differences from Jónsson et al., 2017 (in black). Solid points are statistically significant differences at the 5% level, accounting for multiple tests. Note that the left panel is a duplicate of Fig. 4b, to enable a direct comparison between the results with and without matching for replication timing. The shaded regions in the right panel indicate the range of replication timing (between -0.5 and 0.5) used in this analysis.

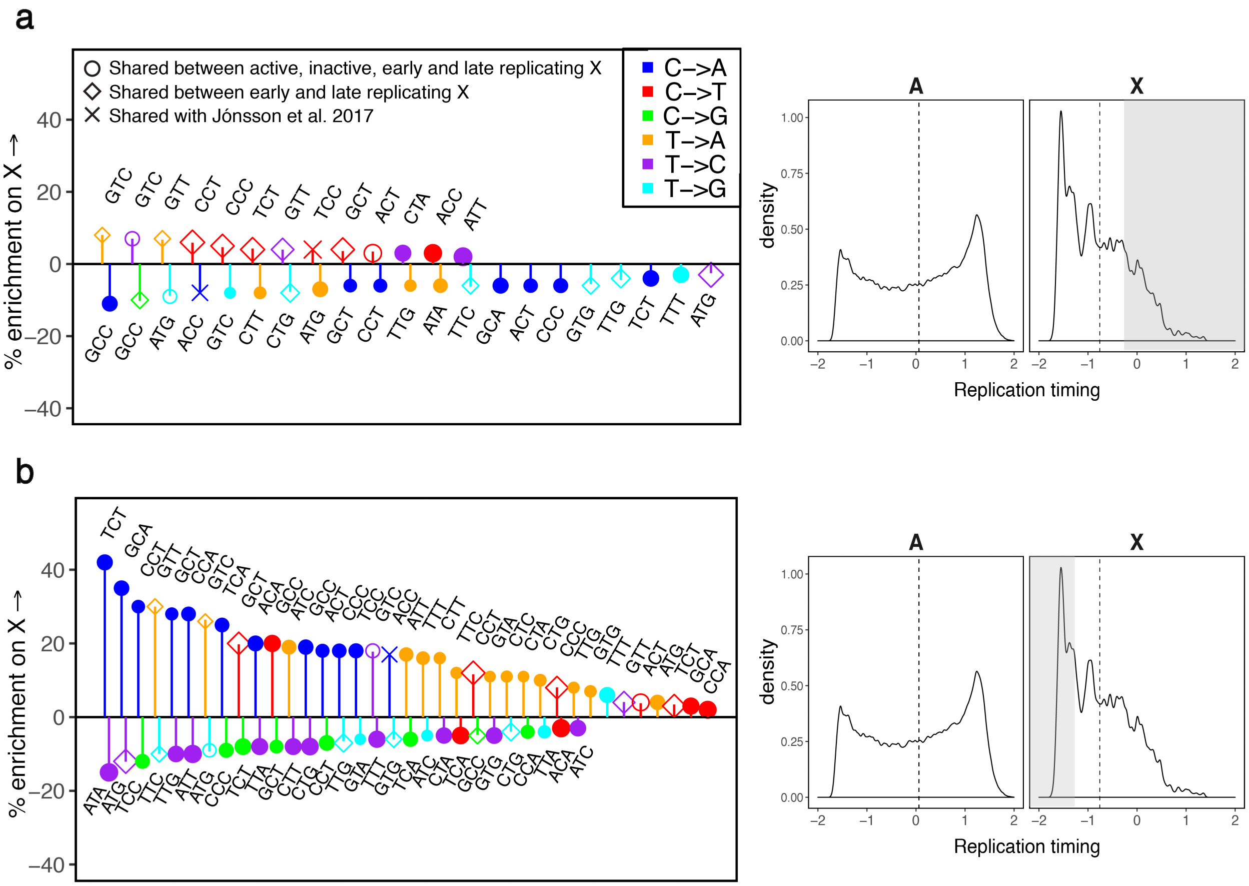


**Fig. S14.** Comparison of the mutation spectrum in early and late replicating compartments of the X chromosome versus autosomes. The pseudoautosomal region (PAR) and CpG sites are excluded from this analysis. Only significant differences are shown. Positive and negative effects are ranked separately in order of effect size from left to right. The size of the marker reflects the number of mutations of that type. Hollow circles indicate the three mutation types that are significantly different in both early and late replicating compartments of the X relative to autosomes and also found to be significant differences in both the escaped and inactive compartments of the X relative to autosomes. Crosses denote mutation types reported as significant sex differences by Jónsson et al., 2017. Hollow diamonds represent other mutation types that are significantly different in both early and late replicating compartments of the X relative to autosomes. The distribution of replication timing for the X and autosomes is shown in the right panels, with the mean replication timing on the X and autosomes represented by dashed vertical black lines. Shaded regions indicate the range of replication timing used in the corresponding analysis. (a) Enrichment of mutation types in early replicating regions of the X chromosome (replication timing score > -0.25) relative to autosomes. (b) Enrichment of mutation types in late replicating regions of the X chromosome (replication timing score < -1.25), relative to autosomes.

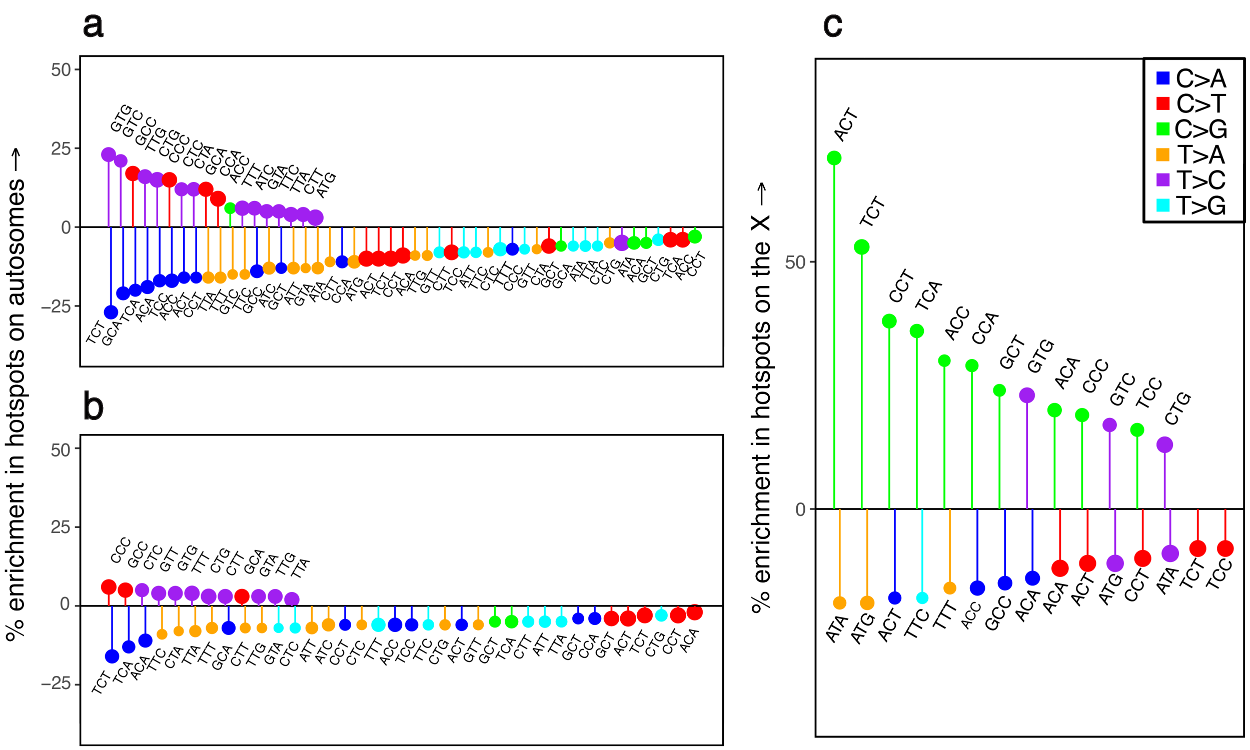


**Fig. S15.** Comparison of the mutation spectrum in regions identified as recombination hotspots, relative to autosomal non-hotspots. These analyses exclude regions on autosomes rich in clustered de novo mutations, identified by Jónsson et al., 2017. CpG sites are excluded. Only significant differences are shown. Positive and negative effects are ranked separately in order of effect size from left to right. The size of the circle reflects the number of mutations of that type. (a) The mutation spectrum in hotspots of DMC1-binding measured on autosomes in males, defined as autosomal regions with hotspot intensity > 0. (b) The mutation spectrum in female crossover hotspots, defined as regions with standardized recombination rate > 10. (c) The mutation spectrum in hotspots of DMC1-binding measured on the X chromosome in males, defined as regions outside the PAR with hotspot intensity > 0.
